# Lcn2 deficiency leads to social impairments independent of maternal immune activation

**DOI:** 10.1101/2025.06.27.661499

**Authors:** Martyna Pekala, Sylwia Zawiślak, Sandra Romanis, Karolina Nader, Aleksandra Cabaj, Anna Madecka, Alicja Puścian, Ewelina Knapska, Robert Pawlak, Leszek Kaczmarek, Katarzyna Kalita

## Abstract

Maternal infection during pregnancy is a well-established risk factor for neurodevelopmental disorders (NDDs), yet the underlying molecular mechanisms remain poorly understood. Lipocalin-2 (Lcn2), an innate immune protein that is highly upregulated during infection, also affects neuronal and glial function. This study investigates the role of Lcn2 in shaping brain development, particularly after maternal immune activation (MIA). To mimic maternal infection, pregnant mice received intraperitoneal injections of either lipopolysaccharide (LPS) or saline on embryonic days 16 to 18 to model infection during the second trimester of pregnancy in humans. We first showed that *Lcn2* mRNA is expressed in the fetal brain and that MIA significantly upregulates *Lcn2* mRNA and protein in the hippocampus and neocortex of both sexes. To assess functional relevance, we employed Lcn2 heterozygous females to generate wild-type and Lcn2 KO offspring from the MIA and control groups. Both female and male offspring underwent a battery of behavioral assays. Lcn2 deletion and MIA independently induced deficits in social behavior and increased repetitive behavior phenotypes relevant to NDDs in adult animals. However, their combination did not exacerbate these effects, suggesting an occlusion effect. Interestingly, no deficits were observed in the learning and memory task. To investigate potential shared molecular mechanisms, we performed RNA sequencing of the fetal forebrain 4 hours after the final LPS injection. This analysis revealed an overlapping group of differentially expressed genes in the Lcn2 KO and MIA groups, indicating convergence on similar transcriptional pathways that may underlie the observed behavioral phenotypes. These results suggest that while Lcn2 may not mediate the pathological effects of prenatal immune challenge, it plays a critical role in normal brain development.

## Introduction

Brain development is a complex and dynamic process that begins early in fetal life and involves the precise formation of neuronal connections essential for cognitive, emotional, and social functioning. A growing body of epidemiological evidence shows that the prenatal period is particularly sensitive to environmental influences, which may disrupt these finely tuned developmental processes and increase the risk of neurodevelopmental disorders (NDDs) (Han et al., 2021b). Among these environmental risk factors, prenatal exposure to maternal infections has emerged as a significant contributor, with numerous systematic reviews and meta-analyses linking it to an increased incidence of autism spectrum disorder (ASD), schizophrenia, and attention-deficit/hyperactivity disorder (ADHD) in offspring (Ayubi and Mansori, 2022; Jiang et al., 2016; Saatci et al., 2021; Tioleco et al., 2021; Zhou et al., 2021; Zhu et al., 2022). Timing appears to be an important factor, with some studies suggesting that infections occurring during the second trimester are particularly associated with a higher risk of NDDs (Jiang et al., 2016; Saatci et al., 2021).

Although the exact mechanisms by which prenatal environmental insults lead to NDDs remain unclear, the maternal immune activation (MIA) hypothesis is one of the most prominent explanations. It proposes that the maternal immune response, characterized by elevated levels of circulating cytokines, rather than the pathogen itself, disrupts fetal brain development (Han et al., 2021a). Supporting this hypothesis are epidemiological studies showing that not only maternal infections, but also chronic inflammatory conditions such as autoimmune diseases, asthma, and obesity, are associated with an increased risk of NDDs in offspring (Han et al., 2021b). Robust experimental evidence also comes from animal models, in which maternal immune activation is induced by such agents as bacterial endotoxin lipopolysaccharide (LPS) or synthetic viral RNA analog Poly(I:C), mimicking bacterial and viral infections, respectively (Meyer et al., 2009). Numerous studies using these models have demonstrated that MIA results in persistent behavioral abnormalities in offspring, accompanied by molecular, anatomical, and functional alterations in the brain (Boksa, 2010; Dunaevsky and Bergdolt, 2019). These behavioral impairments often include deficits in social interaction, increased repetitive behaviors, and cognitive dysfunction, hallmarks resembling the core symptoms of various NDDs (Dunaevsky and Bergdolt, 2019; Hooley, 2010; Leekam et al., 2011; Solek et al., 2018). While such findings strongly support the role of maternal immune signaling in shaping neurodevelopment, the precise molecular mechanisms involved remain poorly defined.

Lipocalin-2 (Lcn2) is an extracellular protein that was initially identified as a part of the innate immune response, released upon infection to bind bacterial siderophores and restrict pathogen growth (Flo et al., 2004; Goetz et al., 2002). Under physiological conditions, Lcn2 levels in the brain are low; however, its expression is markedly upregulated during pathological conditions, including inflammation and following intraperitoneal administration of high doses of LPS (Chia et al., 2011; Hamzic et al., 2013; Ip et al., 2011; Jin et al., 2014; Kang et al., 2018; Marques et al., 2008; Xing et al., 2014; Zhao et al., 2016). Although numerous studies have linked elevated Lcn2 expression to the exacerbation of inflammatory responses, other evidence suggests that Lcn2 may also exert anti-inflammatory effects, highlighting its context-dependent role in immune regulation (Gasterich et al., 2021; Jin et al., 2014; Kang et al., 2018; Kim et al., 2023; Lee et al., 2011; Lee et al., 2009; Li et al., 2023; Vichaya et al., 2019).

Importantly, beyond its immunological functions, Lcn2 is essential for maintaining normal neuronal structure and behavior. Lcn2-knockout (Lcn2 KO) mice exhibit anxiety- and depression-like behaviors, along with deficits in spatial learning (Ferreira et al., 2013; Ferreira et al., 2018; Mucha et al., 2011; Skrzypiec et al., 2013). These behavioral abnormalities are accompanied by impaired adult neurogenesis, altered dendritic arborization, and changes in dendritic spine density. In addition, Lcn2 has been implicated in synaptic plasticity, as in vitro studies show that recombinant Lcn2 reduces membrane expression of NMDA receptor subunits, promotes immature spine morphology, and impairs long-term potentiation (LTP) in hippocampal slices (Doliwa et al., 2024; Kim et al., 2024).

Despite these insights, the function of Lcn2 in the developing brain, specifically its role in modulating prenatal inflammatory responses, remains unexplored. In the present study, we employed a murine MIA model induced by low doses of LPS to address this gap. We demonstrate that MIA upregulates Lcn2 expression in the fetal brain and induces behavioral abnormalities reminiscent of symptoms associated with NDDs. Strikingly, Lcn2 deletion alone produced comparable behavioral deficits, and combining MIA with Lcn2 deficiency failed to exacerbate these effects, suggesting an occlusion phenomenon. To uncover potential common molecular pathways, we performed bulk RNA sequencing on fetal forebrain tissue and identified a shared set of differentially expressed genes in both Lcn2 KO and MIA groups. These results suggests that, while Lcn2 may not directly mediate the deleterious impact of prenatal immune challenge, it plays a crucial role in normal brain development.

## Materials and methods

### Animals

Wild-type pregnant C57BL/6J and transgenic pregnant females with a knockout of the Lcn2 gene (strain: B6.129P2-Lcn2tm1Aade/AkiJ, stock #024630, The Jackson Laboratory, USA), maintained on a C57BL/6J genetic background, were used in the study (Flo et al., 2004; Mucha et al., 2011). All animals were bred and housed at the Animal Facility of the Nencki Institute under standard laboratory conditions: a 12-h light/dark cycle, ambient temperature maintained at 20–23°C, relative humidity between 45–65%, and 10–15 air changes per hour. Animals had ad libitum access to food and water. Mice were housed in individually ventilated cages (391 × 199 × 160 mm) in groups of 4–5 animals. Pregnant females were separated around gestational day 14 and housed individually until weaning of the offspring. For behavioral testing, males and females were kept in larger cages (395 × 346 × 213 mm) in groups of 9–14 animals. All cages were enriched with poplar wood bedding and nesting materials (wooden blocks, cellulose cotton rolls, and wood wool). All experimental procedures were performed following the European Communities Council Directive of 24 November 1986 (86/609/EEC), the Animal Protection Act of Poland, and approved by the 1st Local Ethics Committee in Warsaw (approval no. 698/2018).

### Maternal immune activation (MIA) model

A maternal immune activation model was employed to induce systemic immune responses in pregnant mice using intraperitoneal injections of lipopolysaccharide, a component of the outer membrane of Gram-negative bacteria. Female mice aged 10–35 weeks were mated with a male overnight. C57BL/6J females were mated with C57BL/6J males, while Lcn2 heterozygous females were mated with either wild-type or knockout males. The following morning, body weight was recorded, and the presence of vaginal plugs was checked to confirm mating. Pregnancy was further verified based on subsequent weight gain. Beginning on gestational day 16, pregnant females received daily intraperitoneal injections of LPS derived from *Escherichia coli* (serotype O111:B4, Sigma-Aldrich, cat. no. L4391-1MG) dissolved in sterile saline (Injectio Natrii Chlorati Isotonica, Polpharma, Poland) at a concentration of 5 μg/mL. The dosage was 40 μg/kg of body weight, administered in a volume appropriate for each animal for three consecutive days. Control animals received equivalent volumes of sterile saline. Offspring at various developmental stages (prenatal and postnatal) were used for subsequent analyses.

### Sex determination of mouse fetuses by polymerase chain reaction (PCR)

Genomic DNA was isolated from the mice’s tails to determine the sex of the animals. The tissue was incubated in lysis buffer (100 mM NaCl, 100 mM Tris-HCl, 0.5% SDS, 1 mM EDTA, proteinase K 1:10) overnight at 55 °C. The next day, proteinase K was inactivated by 10 min incubation at 98 °C. The obtained lysate was then diluted in deionized water in a ratio of 1:40 and used as template DNA for amplification using primers 5’-CACCTTAAGAACAAGCCAATACA-3’ and 5’-GGCTTGTCCTGAAAACATTTGG-3 (Tunster, 2017).

### Measurement of Lipocalin-2 Concentration in Maternal Plasma

Lcn2 levels were quantified in maternal plasma 24 hours after the final LPS injection using the Mouse Lipocalin-2/NGAL DuoSet ELISA kit (# DY1857, R&D Systems, Minneapolis, MN, USA), following the manufacturer’s instructions. Blood was collected via cardiac puncture using a 1 ml syringe and transferred to EDTA-coated tubes to prevent coagulation. Samples were centrifuged within 30 minutes of collection at 4 °C for 15 minutes at 1000 × g. The plasma fraction was carefully separated and subjected to a second centrifugation at 4 °C for 10 minutes at 10,000 × g to remove any remaining cellular debris. The plasma was aliquoted and stored at −80 °C until analysis.

### Western blot analysis

Sixty micrograms of protein extracts collected from the cortex or hippocampus of the E19 fetuses were run on polyacrylamide gels under reducing conditions. The standard Western blot procedure was performed using anti-Lcn2 (#AF 3508, R&D Systems, Minneapolis, MN, USA). To monitor equal total protein levels, the blots were re-probed with β-actin (#A1978, Sigma-Aldrich, Saint Louis, MO, USA) antibodies. The chemiluminescent method was used for signal detection. To quantify individual bands, a scan of X-ray films was analyzed by densitometry using ImageJ.

### RNA preparation and quantitative real-time PCR

Total RNA was isolated from the fetal brain using the RNeasy Mini Kit (#74104, Qiagen, Hilden, Germany). RNA was reverse transcribed with SuperScript™ IV VILO (# 11766050, Thermo Fisher Scientific, Waltham, MA, USA) according to the manufacturer’s instructions. cDNA was amplified with TaqMan probes (Lcn2 Mm01324470_m1, Gapdh Mm99999915_g1, En2 Mm00438710_m1, Pax2: Mm01217939_m1, ThermoFisher, Waltham, MA, USA) that were specific for the mouse. Quantitative real-time PCR was performed using TaqMan Fast Advanced Master Mix (#4444556, Thermo Fisher Scientific, Waltham, MA, USA) with StepOne Real-Time PCR System (Thermo Fisher Scientific, MA USA). Fold changes in expression were determined using the ΔΔCT relative quantification method. The values were normalized to relative amounts of Gapdh.

### Eco-HAB® testing of social behaviors

Eco-HAB® is a fully automated, RFID-based, open-source system designed to assess naturalistic social behaviors in group-housed mice (Puścian et al., 2016). The apparatus consists of four interconnected polycarbonate compartments (30 × 30 × 18 cm), linked by tubular corridors (internal diameter: 36 mm; external diameter: 40 mm). Mice implanted with subcutaneous RFID microchips were automatically tracked by antennas placed in the corridors. Behavioral data were recorded continuously and analyzed using the Python-based pyEcoHAB library (https://github.com/Neuroinflab/pyEcoHAB).

Mice (8-11 weeks) were tested in the Eco-HAB® system over six consecutive days, following a three-phase protocol: adaptation (days 1–2), sociability assessment (days 3–5), and olfactory stimulus presentation (day 6). At the onset of the dark phase on day 6, soiled bedding from unfamiliar, age- and sex-matched wild-type mice (same strain) was placed behind a perforated partition in one compartment to serve as a social olfactory cue. Clean bedding was used in the opposite compartment as a neutral control.

To characterize spontaneous social interactions within the group, in-cohort sociability was calculated as the total time each pair of mice spent together minus the time they would be expected to spend together based on their individual preferences for occupying specific compartments within the Eco-HAB® system (Puścian et al., 2016; Roszkowska et al., 2022). This parameter was calculated for every pair of animals within each experimental group during dark phases 3, 4, and 5. Additionally, the approach to social odor was evaluated by calculating the increase in the proportion of time each animal spent in the compartment containing a novel social odor compared to the time spent in the compartment with a neutral olfactory cue relative to the corresponding period during the preceding dark phase.

### Three-chamber test

The three-chamber test was used to assess social behavior deficits in mice. Testing was conducted in a non-transparent box measuring 63 × 44 × 25 cm, divided into three equal compartments (21 × 44 × 25 cm) by walls containing openings that allowed the animal free access to all chambers. In the center of each of the two outer chambers, an inverted cylindrical metal wire cup (15 cm in height, 12 cm in diameter) was placed to serve as a stimulus container. Mice were tested individually in two consecutive 10-minute phases. During the first phase (habituation), both wire cups were empty, and the mouse was placed in the center chamber with free access to all three compartments to explore the apparatus. In the second phase, one wire cup contained a control olfactory stimulus (clean bedding), while the other contained a social olfactory stimulus (bedding soiled by unfamiliar, age- and sex-matched wild-type conspecifics of the same strain). Behavior was recorded using a video camera connected to a computer and analyzed with EthoVision XT software (Noldus Information Technology, Wageningen, Netherlands), which enables automated tracking of animal movement and interaction. The primary behavioral parameter was the duration of interaction with each wire cup, defined as sniffing, close investigation, or climbing. A social odor preference index (SI) was calculated using the formula: SI = (TS − TNS) / (TS + TNS), where *TS* represents the time spent interacting with the cup containing the social stimulus, and *TNS* represents the time spent interacting with the cup containing the control stimulus.

### Marble burying test

The marble burying test was conducted in a clean, transparent plastic cage (27 × 21 × 14 cm) filled with approximately 5 cm of tamped-down bedding to create an even, flat surface. The cage was covered with a second transparent plastic cage of the same dimensions. Twelve identical glass marbles (15 mm in diameter) were arranged in a regular pattern across two rows on the bedding surface. Each mouse was placed individually into the experimental cage and allowed to explore for 15 minutes. After each session, the mouse was returned to its home cage, and the number of marbles buried, defined as being covered by bedding to at least two-thirds of their depth, was recorded as a measure of repetitive digging behavior.

### Autistic-like score

Autistic-like score is a composite behavioral index designed to quantify autism-like traits, specifically impairments in social interaction, reduced interest in social stimuli, and increased repetitive behaviors. The score was computed based on previously described methods (Bosch-Bouju et al., 2016; Hilal et al., 2024), using the standardized values of four behavioral parameters: sociability (EcoHAB® test), approach to social odor (EcoHAB® test), social odor preference index (three-chamber test), and number of marbles buried (marble burying test). For each parameter, standardization was performed as follows: (x – min value) / (max value – min value), where *x* is the raw value for a given animal, and *min* and *max* are the minimum and maximum values across the entire cohort. Given that approach to social odor and social odor preference index assess overlapping behavioral dimensions, their standardized values were averaged so that each of the three behavioral domains (social interaction, social interest, and repetitive behavior) contributed equally to the final score. To ensure consistent directionality of the score (i.e., higher values reflecting stronger autistic-like behavior), the standardized scores for social interaction and social interest were inverted using 1-standardized score. The autistic score was then calculated as: (1-score _social interaction_) + (1 – score _social interest_) + score _repetitive behavior_. This procedure yields a score ranging from 0 to 3, with higher values indicating more pronounced autistic-like traits.

### IntelliCage system for testing reward-motivated learning

IntelliCage (TSE Systems, Berlin, Germany) is an automated system for monitoring behavior in group-housed mice (Knapska et al., 2013; Knapska et al., 2006). It consists of a home cage (55 × 37.5 × 20.5 cm) with four experimental corners, each equipped with sensors and controlled via IntelliCagePlus software (v3.3.7.0). Access to each corner is limited to one mouse via a 3 cm tube. An overhead sensor detects presence, while an RFID antenna identifies individual mice via implanted transponders. Each corner contains two water bottles behind motorized doors, which open following a nose-poke.

To establish a preference for a positively valenced stimulus, mice were trained to seek a 10% sucrose solution in the IntelliCage system. A two-day adaptation period followed, with bottle-access doors open and free access to all corners. Next, doors were closed for two days, and mice learned to nose-poke for water access. Doors opened for 5 seconds following a correct response. At the end of this session, the least preferred corner (i.e., the one with the fewest visits) was identified for each mouse. In the following three-day phase, water access was restricted to the individually least preferred corner. Instrumental responses to both bottles in that corner were recorded. The bottle associated with fewer responses was then replaced with a 10% sucrose solution, available for the next five days. Animal activity, learning to discriminate between bottles containing plain and sucrose-sweetened water, and sucrose preference were assessed.

### RNA Sequencing

Tissue from an E18 mouse forebrain was quickly dissected and stored in RNAlater solution (Invitrogen, AM7020) at 4°C for 24 hrs, supernatant was removed and tissue kept kept at −70°C for further use. Total RNA was isolated using the RNeasy Mini Kit (#74104, Qiagen, CA, USA) according to the manufacturer’s protocol. Quality and integrity of total RNA were assessed with Agilent 2100 Bioanalyzer using an RNA 6000 Nano Kit (#5067-1511, Agilent Technologies, Ltd., CA, USA) In total, strand-specific polyA-enriched RNA libraries were prepared using the KAPA Stranded mRNA Sample Preparation Kit according to the manufacturer’s protocol (#07962207001, Kapa Biosystems, MA, USA). Briefly, mRNA molecules were enriched from 500ng of total RNA using poly-T oligo-attached magnetic beads (#07962207001, Kapa Biosystems, MA, USA). The obtained mRNA was fragmented, and the first-strand cDNA was synthesized using a reverse transcriptase. Second, cDNA synthesis was performed to generate double-stranded cDNA (dsDNA). Adenosines were added to the 3′ ends of dsDNA, and adapters were ligated (#E7600S, adapters from NEB, Ipswich, MA, USA). Following the adapter ligation, uracil in a loop structure of the adapter was digested by USER enzyme from NEB (#E7600S, Ipswich, MA, USA). Adapters containing DNA fragments were amplified by PCR using NEB starters (#E7600S, Ipswich MA, USA). Library evaluation was done with Agilent 2100 Bioanalyzer using the Agilent DNA High Sensitivity chip (#5067-4626, Agilent Technologies, Ltd. CA, USA). The mean library size was 300bp. Libraries were quantified using a Quantus fluorometer and QuantiFluor double-stranded DNA System (#E4871, Promega, Madison, Wisconsin, USA). Libraries were paired-end sequenced (2 × 151bp) on NovaSeq 6000 (Illumina, San Diego, CA, USA). RNA sequencing was performed in the Laboratory of Sequencing, Nencki Institute of Experimental Biology, Poland.

### Data preprocessing and analysis

Quality control of reads was performed using FastQC (v. 0.11.9). Adapter and quality trimming were performed using cutadapt (v. 3.4) and TrimGalore (v. 0.6.7), respectively. TrimGalore quality parameter was set to 25. Alignment of reads to the murine reference genome (GRCm39) was performed using STAR (v. 2.7.9a) with default settings. The GTF annotation file was used from the 105 Ensembl release. Duplicate reads were marked using Picard MarkDuplicates (v. 2.27.4-SNAPSHOT). Final quality control was collated with MultiQC (v. 1.13) from RSeQC (v. 3.0.1) and the tools described above. Reads were summarized and counted by featureCounts (v. 2.0.0) on paired-end reads, with only primary alignments and reversely stranded reads counted. The minimum mapping quality score required for a read to be counted was set to 3. The differential analysis was performed using DESeq2 (v. 1.34) with default parameters. If the sum of counts for a specific feature was less than 10 in all samples, this feature was discarded from the analysis. Plots were generated using ComplexHeatmap (v. 2.10.0) packages, GraphPad and overlapping genes were identified using Venn diagram tool (http://bioinformatics.psb.ugent.be/webtools/Venn/), followed by the exact hypergeometric probability significance test of the overlap using a web-based program (http://nemates.org/MA/progs/overlap_stats.html)(Gao et al., 2024). Significantly differentially expressed genes (DEGs) (FDR-adjusted p-value < 0.05) obtained from the RNA sequencing were used as input for the pathway analysis. The functional analysis was performed using Metascape https://www.metascape.org/ (Zhou et al., 2019). The function of DEGs was investigated using Gene Oncology (GO) of biological processes (BP). For identified DEGs, function and pathway enrichment analysis were carried out using the following ontology sources: Gene Ontology and KEGG Pathway with a p < 0.01, a minimum count of 3, and an enrichment factor >1.5 were collected and grouped into clusters.

### Statistical analyses

Sample sizes and detailed statistical information are provided in the text, figures, or their respective legends. Normality of data distribution was assessed using the D’Agostino-Pearson test. Data with normal distribution were analyzed using Student’s *t*-test, one-way ANOVA, two-way ANOVA, two-way repeated-measures ANOVA, or three-way repeated-measures ANOVA, followed by Šídák’s, Tukey’s, or Fisher’s LSD post hoc tests for multiple comparisons. For comparisons involving more than two groups with non-normal distributions, logarithmic transformation was applied (Y = Ln (Y), or Y = Ln (Y + 1) for data containing zero values). For two-group comparisons, the Mann–Whitney *U* test was applied. Fetal survival data were analyzed using the chi-square (χ²) test. Differences between experimental groups were considered statistically significant at *p* < 0.05. All analyses were performed using GraphPad Prism (version 8.0.2, GraphPad Software).

## Results

### Temporal expression of Lipocalin-2 in the developing hippocampus and its sensitivity to maternal immune activation

While the role of Lcn2 in the adult brain has been extensively studied (Ferreira et al., 2013; Ip et al., 2011; Mucha et al., 2011; Yan et al., 2024), its function during brain development remains poorly understood. To determine whether *Lcn2* mRNA in the brain is developmentally regulated, we performed RT-qPCR using RNA isolated from the hippocampi of wild-type C57BL/6J mice of both sexes across various prenatal and postnatal stages, starting from embryonic day 16 (E16) to adult mice (two months old). *Lcn2* mRNA expression was detectable in the hippocampus as early as the prenatal period and remained present at all examined developmental stages (Figure 1A). The lowest expression levels were observed at E16. A significant increase in Lcn2 expression at consecutive stages of development was observed (E16 vs. E19 p = 0.0182; E16 vs. P0 p < 0.0001; E16 vs. P7 p = 0.0068; E16 vs. P14 p < 0.0001; E16 vs. P21 p = 0.0002; E16 vs. 5 weeks p = 0.0004; E16 vs. adult p = 0.0080; Figure 1A). *Lcn2* mRNA expression peaked at birth (P0), although statistical differences were found only in comparisons with E16 (P0 vs. E16 p < 0.0001; Figure 1A), E19 (P0 vs. E19 p = 0.0036; not shown on graph), P7 (P0 vs. P7 p = 0.0106; not shown on graph), and adult mice (P0 vs. adult p = 0.0177; not shown on graph). During the postnatal period, Lcn2 expression remained relatively stable into adulthood.

**Fig. 1.**
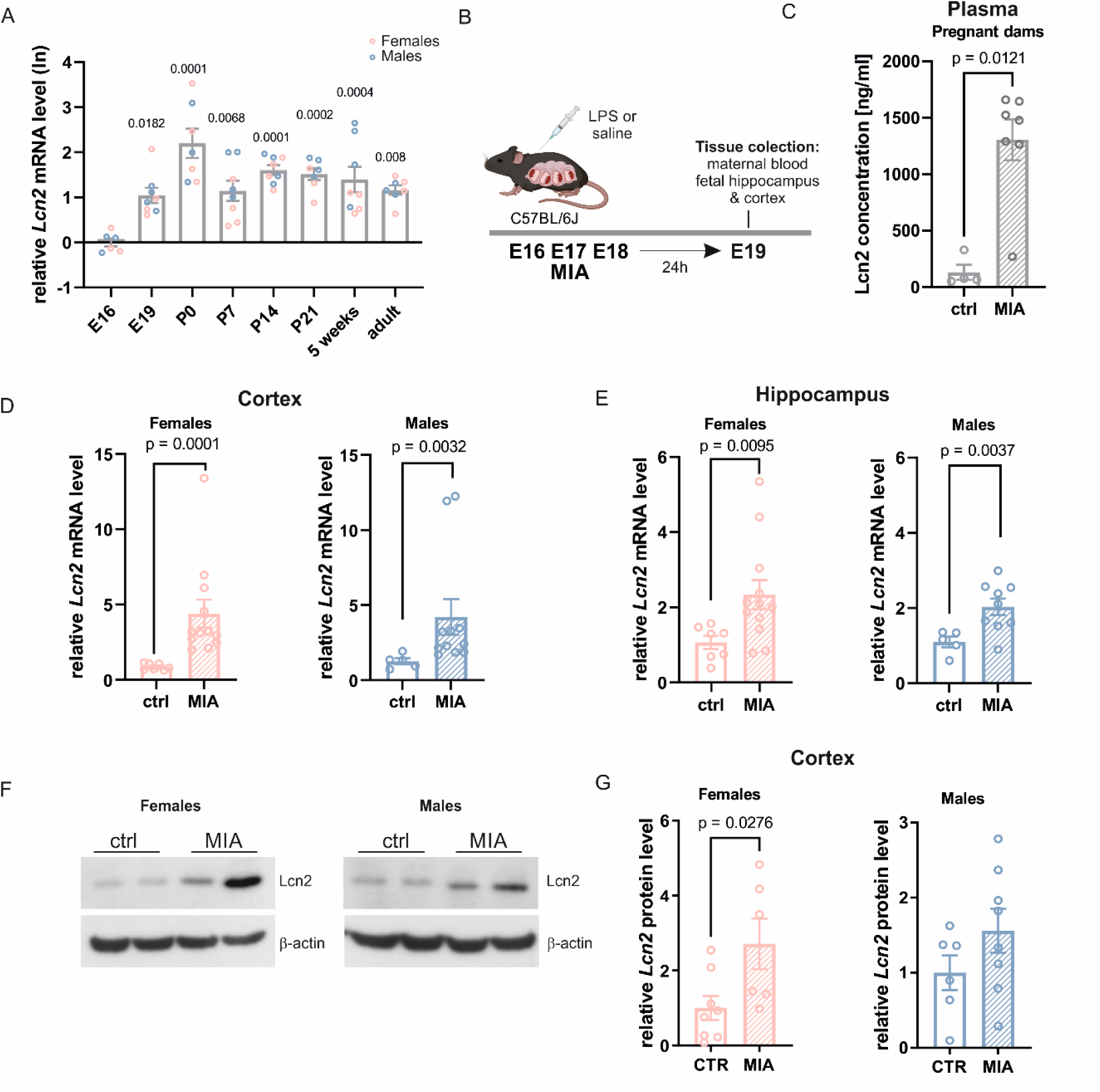
Lipocalin-2 is induced in the offspring brain following maternal immune activation induced by LPS. **A.** *Lcn2* mRNA expression in the hippocampus during pre- and postnatal development in mice. E16 n = 6, E19 n = 8, P0 n = 7, P7 n = 8, P14 n = 8, P21 n = 7, 5 weeks n = 8, adulthood n = 7 (offspring sex is marked with colors). Data were log-transformed (Y = Ln(Y)) before analysis. One-way ANOVA (F (7, 51) = 8.517, p < 0.0001) and Tukey’s post hoc test. **B.** Scheme of the experiment. LPS (40 µg/kg) was administered to pregnant C57BL/6J6 mice on embryonic days E16, E17, and E18. At E19, samples were collected for further analysis. **C.** Lcn2 protein level in the plasma from pregnant dams at 24 hours after the last LPS injection, measured by enzyme-linked immunosorbent assay (ELISA). ctrl n = 4, MIA n = 7. Mann-Whitney test analysis. **D, E.** *Lcn2* mRNA expression levels in the cerebral cortex and hippocampus of female and male fetuses, measured 24 hours after the final injection of LPS or saline in pregnant dams, females: ctrl n = 7, MIA n = 12; males: ctrl n = 5, MIA n = 11 or 9. Mann-Whitney test analysis. **F.** Representative immunoblot of Lcn2 in the cortex and hippocampus from female and male offspring 24 hours after the last LPS injection, **G.** Quantification of immunoblot for Lcn2 band intensity, females: ctrl n = 8, MIA n = 6; males: ctrl n = 6, MIA n = 8. Student’s t-test analysis. All data are presented as mean ± SEM.

To examine how maternal infections affect brain development and whether Lcn2 plays a role in this process, we employed the MIA model. In this approach, pregnant mice were administered bacterial endotoxin to mimic prenatal infection. Beginning on E16, C57BL/6J dams received daily intraperitoneal injections of LPS at a dosage of 40 μg/kg body weight for three consecutive days or sterile saline (Fig. 1B). To assess the MIA model’s impact on pregnancy progression, we tracked body weight gain in pregnant females (Supplementary Fig. 1A). Pregnant dams were weighed every morning from the day preceding the first injection through the final injection day (E18). We calculated the percent weight gain based on the initial weight recorded on E15. A significant decrease in maternal weight gain was found after the first LPS injection compared to control females receiving saline (after the first dose – E17, ctrl vs. MIA: p = 0.0068; Supplementary Fig. 1B). This effect was temporary, as no significant difference was observed after the second injection (after second dose – E18, ctrl vs. MIA). Additionally, LPS injections notably affected fetal survival, with nearly 50% of pregnancies in the LPS-treated group resulting in stillbirths (χ² test, ctrl vs. MIA: p = 0.0058; Supplementary Fig. 1C).

To assess how the MIA model affects Lcn2 levels during pregnancy, we collected serum from females 24 hours after the last LPS injection. The level of Lcn2 was evaluated using the ELISA method. Pregnant dams after MIA exhibited significantly increased Lcn2 levels in the blood (ctrl vs. MIA, p = 0.0121, Fig. 1C). Given that we demonstrated the presence of *Lcn2* mRNA in the developing hippocampus, we aimed to determine whether its expression is sensitive to maternal immune activation. To this end, RT-qPCR was performed using RNA isolated from the cortex and hippocampi of male and female fetuses collected from wild-type C57BL/6J dams 24 hours after the final injection of either saline or LPS. Maternal immune activation resulted in a significant upregulation of *Lcn2* mRNA expression in both the cortex (Fig. 1D) and hippocampus (Fig. 1E) of fetuses of both sexes compared to controls (cortex: females, p < 0.0001, males, p = 0.0032; hippocampus: females, p = 0.0095, males, p = 0.0037). Notably, the average increase in *Lcn2* mRNA expression was greater in the cortex (females: control 0.87 ± 0.08 vs. MIA 4.39 ± 0.93; males: control 1.26 ± 0.20 vs. MIA 4.22 ± 1.18) than in the hippocampus (females: control 1.07 ± 0.17 vs. MIA 2.34 ± 0.39; males: control 1.09 ± 0.14 vs. MIA 2.03 ± 0.22). To confirm these results at the protein level, we performed Western blot analysis on samples isolated from fetuses’ brains of both sexes to measure Lcn2 levels. We showed a statistically significant upregulation of Lcn2 protein levels in the female cortex 24 hours after the last LPS injection (Fig. 1D, F; p = 0.0276). Although a similar increase was observed in males, this result did not reach statistical significance.

### Lcn2 deletion mimics behavioral deficits induced by MIA

After establishing a validated model of MIA characterized by the upregulation of Lcn2 levels in the fetal brain, we examined how maternal infection affects offspring behavior. To assess the potential role of Lcn2 in MIA-induced behavioral changes, we used transgenic animals. Pregnant Lcn2 Het females received daily intraperitoneal injections of LPS (40 μg/kg) or saline from E16 to E18. Using heterozygous Lcn2 females allowed us to control for maternal genotype-dependent effects on offspring outcomes. After weaning, the offspring from various litters were housed together based on treatment, genotype, and sex. Adult Lcn2 KO and wild-type (WT) offspring of both sexes were later evaluated for behaviors associated with NDDs, such as social interaction, repetitive behavior, and cognitive function. To determine how MIA influences pregnancy progression in Lcn2 heterozygotes, we analyzed body weight changes in pregnant females administered either saline or LPS injections (Supplementary Fig. 2A). MIA induced by LPS resulted in a significant decrease in maternal weight gain after both the first and second injections compared to the saline-treated controls (after first injection – E17, ctrl vs. MIA: p < 0.0001; after second injection – E18, ctrl vs. MIA: p < 0.0001; Supplementary Fig. 2B). LPS exposure also significantly affected fetal viability, with approximately 50% of pregnancies in the MIA group ending in stillbirths (χ² test, control vs. MIA: p = 0.0002; Supplementary Fig. 2C).

To assess spontaneous and stimulus-driven social behaviors, we used Eco-HAB®, an automated RFID-based system that continuously monitors social behavior in group-housed mice living under seminaturalistic conditions (Puścian et al., 2016; Roszkowska et al., 2022). Eco-HAB® consists of four chambers connected by tube-like corridors; two chambers provide access to food and water, while the other two are equipped with transparent, perforated separators that allow for the presentation of olfactory stimuli without direct physical contact (Fig. 2B). Following a two-day habituation period, social interactions among all pairs of mice were monitored from days 3 to 5 (Fig. 2C). We quantified the in-cohort sociability parameter, which is defined as the time mice voluntarily spent with other group members. We found that MIA significantly reduced social interactions in WT offspring of both sexes (Fig. 2D, G, E, and H). Notably, deletion of the *Lcn2* gene in saline-treated mice led to a similar decrease in social behaviour - both male and female KO mice exhibited reduced in-cohort sociability compared to WT controls. Interestingly, MIA did not exacerbate this deficit in Lcn2 KO mice (Fig. 2D, G).

**Fig. 2.**
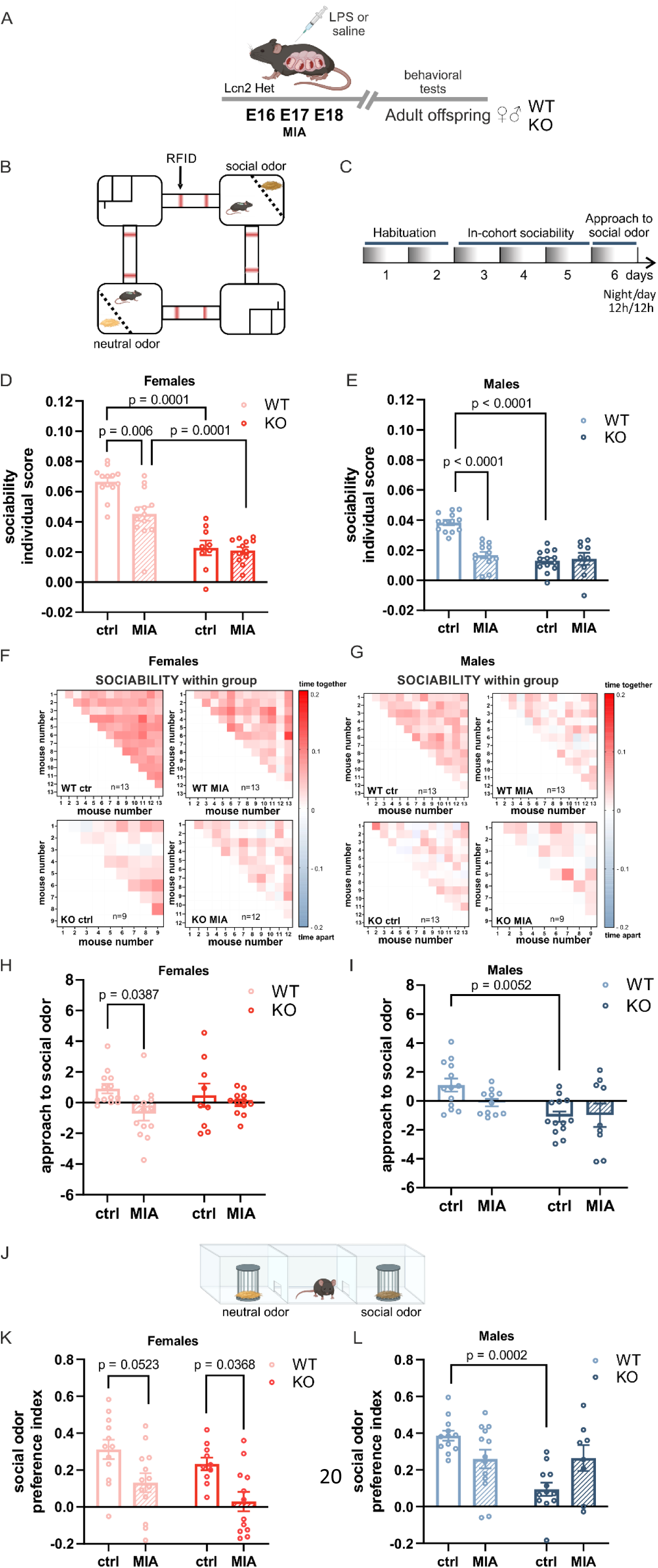
Maternal immune activation or the Lcn2 deletion alters sociability. **A.** Scheme of the experiment. LPS was administered to pregnant Lcn2 Het mice on embryonic days E16, E17, and E18. Behavioral testing was performed in adult offspring. The same cohort of animals was used across all behavioral assessments. **B.** Scheme of the Eco-HAB® apparatus. **C.** Timeline of the experiment in the Eco-HAB® system. **D, E.** Individual score of sociability of females **(D)** and males **(E)** presented as mean value for a given mouse in a group. **(D)** Two-way ANOVA effects of procedure: F(1,43) = 9.429, p = 0.0037, genotype: F(1,43) = 82.52, p < 0.0001, and interaction: F(1,43) = 6.795, p = 0.0125, followed by Tukey’s post hoc test. **(E)** Two-way ANOVA effects of procedure: (F(1,44) = 18.05, p = 0.0001); genotype: F(1,44) = 34.34, p < 0.0001; interaction: F(1,44) = 23.09, p < 0.0001), followed by Tukey’s post hoc test. **F, G.** Raw data from sociability presented as a matrix, where each square represents the time each pair of mice voluntarily spent together. Color intensity corresponds to the strength of their interaction, following the provided scale. **H, I.** Approach to social odor of female **(H)** and male **(I)** mice. **(H)** Two-way ANOVA effect of procedure: F (1, 43) = 5.782; p = 0.0206), genotype: F (1, 43) = 5.782; p = 0.0206, and interaction: F (1, 43) = 1.666; p = 0.2036), followed by Tukey’s post hoc test. **(I)** Two-way ANOVA effect of procedure: F (1, 43) = 0.3486; p = 0.5580, genotype: F (1, 43) = 11.43, p = 0.0015, and interaction: F (1, 43) = 0.6216; p = 0.4348, followed by Tukey’s post hoc test. Females WT ctrl n = 13, WT MIA n = 13, KO ctrl n = 9, KO MIA n = 12; males WT ctrl n = 13, WT MIA n = 13, KO ctrl n = 13, KO MIA n = 9. Due to the non-normal distribution of the data, a logarithmic transformation (Y = Ln[Y]) was applied. Data are presented as mean ± SEM. **J.** Scheme of the three-chamber social approach task. **K, L.** Social odor preference index in females (**K**) and males (**L**). **(K)** Two-way ANOVA effect of procedure: F (1, 43) = 14.88; p = 0.0004, genotype: F (1, 43) = 3.273; p = 0.0774, and interaction: F (1, 43) = 0.05483, p = 0.8160 followed by Tukey’s post hoc test. **(L)** Two-way ANOVA effect of procedure: F (1, 41) = 0.2154, p = 0.6450, genotype: F (1, 41) = 9.547; p = 0.0036, and interaction: F (1, 41) = 10.18; p = 0.0027, followed by Tukey’s post hoc test. Females: WT ctrl n = 12, WT MIA n = 13, KO ctrl n = 10, KO MIA n = 12; males: WT ctrl n = 12, WT MIA n = 13, KO ctrl n = 12, KO MIA n = 8. Data are presented as mean ± SEM.

We also evaluated the approach to social odor, specifically, the preference for novel social cues. This parameter was quantified as the proportion of time spent investigating social (bedding from unfamiliar mice) versus non-social (clean bedding) odors presented behind the perforated separators in the opposing Eco-HAB® compartments on day 6 of the experiment. In WT females, prenatal LPS exposure decreased interest in the social stimulus compared to vehicle-treated controls (WT ctrl vs. WT MIA: p = 0.0387, Fig. 2H). A similar trend was noted in males; however, it did not reach statistical significance (p = 0.25). Conversely, deletion of the *Lcn2* gene diminished social odor preference only in vehicle-treated males (WT ctrl vs. KO control: *p* = 0.0052, Fig. 2I).

To independently validate the findings of Eco-HAB® using a classical and well-established method, we assessed social behavior in the three-chamber test (Fig. 2J). This apparatus evaluates the preference for approaching social and non-social stimuli. To ensure consistency with the Eco-HAB® paradigm, we used clean bedding as the control stimulus, while bedding from unfamiliar mice served as the social stimulus, which was presented under wire cups. We recorded the time the mouse spent interacting with each cup, and a social odor preference index was calculated. Consistent with the Eco-HAB® results, both MIA and *Lcn2* gene deletion reduced the preference for social odor in a sex-dependent manner (Fig. 2K, L). In females, we observed a decreased interest in the social olfactory stimulus following prenatal LPS injections in wild-type mice, with the difference approaching statistical significance (WT ctrl vs. WT MIA: p = 0.0523; Fig. 2K). A similar effect of MIA was noted in female knockout mice (KO ctrl vs. KO MIA: p = 0.0368; Fig. 2K). In males, the three-chamber results aligned with those from the Eco-HAB® test, demonstrating that social odor preference was reduced exclusively in control males lacking the *Lcn2* gene (WT ctrl vs. KO ctrl: p = 0.0002; Fig. 2L).

Next, we evaluated the impact of maternal immune activation and *Lcn2* gene deletion on repetitive behaviors, one of the hallmark features of ASD, by employing the marble-burying test (Fig. 3A). Mice naturally show digging and burying behaviors; however, an increase in these actions is often utilized as a measure for repetitive behavior (Chang et al., 2017; Thomas et al., 2009). Both female and male control mice lacking the *Lcn2* gene buried significantly more marbles than their wild-type counterparts (WT ctrl vs. KO ctrl: females, p = 0.0107, Fig. 3B; males, p = 0.0002, Fig. 3C). Furthermore, in males, prenatal LPS exposure resulted in a higher number of buried marbles in wild-type offspring compared to controls (WT ctrl vs. WT MIA: p = 0.0048; Fig. 3C), while a similar trend was observed in females, although it did not reach statistical significance (p = 0.11; Fig. 3B).

**Fig. 3.**
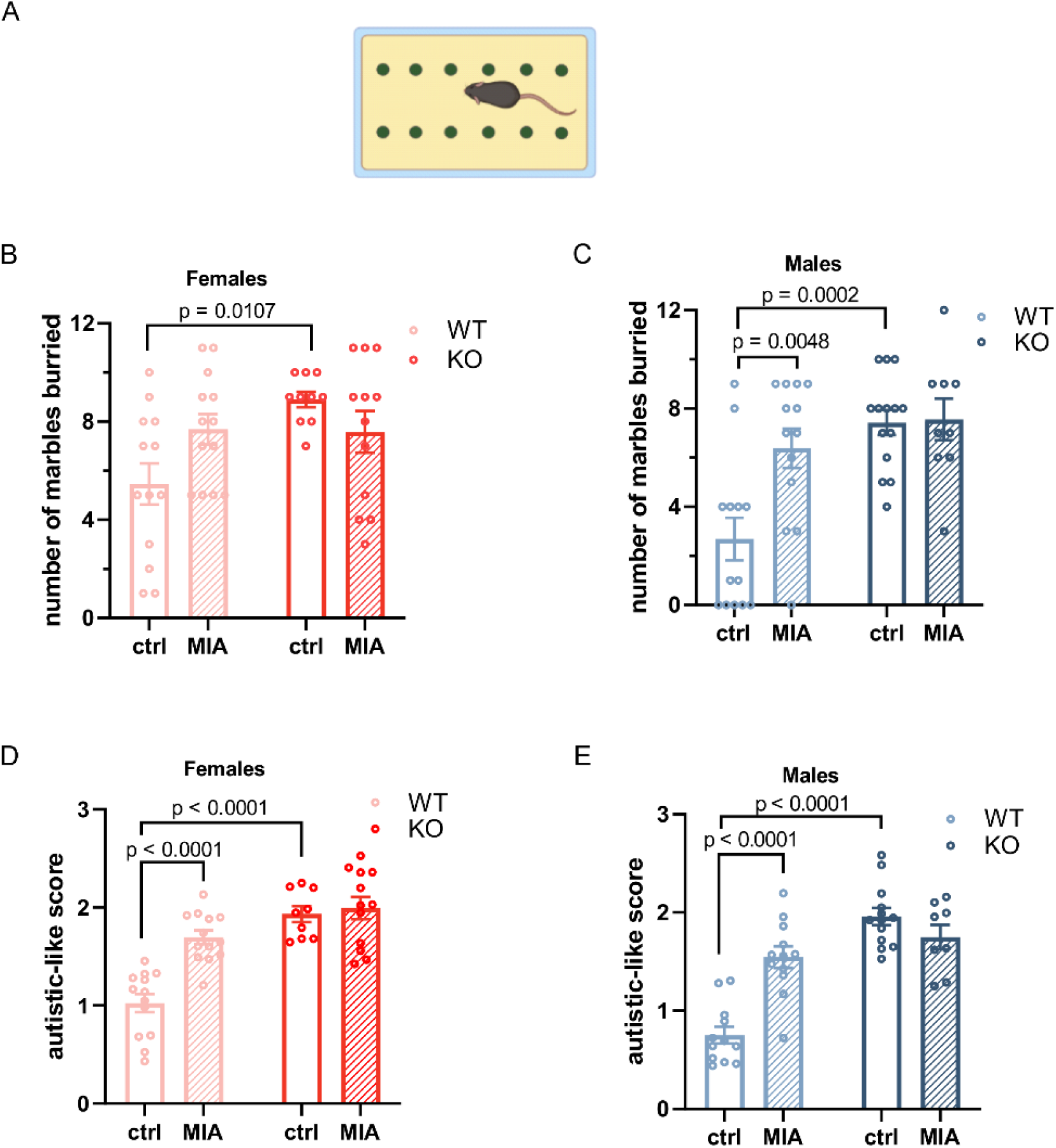
MIA does not exacerbate core autistic-like behaviors in Lcn2 KO mice. **A.** Scheme of the marble burying test. **B, C.** Number of buried marbles in females **(B)** and males **(C)**. (**B**) Two-way ANOVA procedure: F (1, 44) = 0.3982; p = 0.5313, genotype: F (1, 44) = 5.283, p = 0.0263, and interaction: F (1, 44) = 5.997; p = 0,0184, followed by Tukey’s post hoc test. **(C)** Two-way ANOVA procedure: F (1, 45) = 6.185; p = 0.0167, genotype: F (1, 45) = 14.80; p = 0.0004, interaction: F (1, 45) = 5.390; p = 0.0248, followed by Tukey’s post hoc test. Females: WT ctrl n = 13, WT MIA n = 13, KO ctrl n = 10, KO MIA n = 12; males: WT ctrl n = 13, WT MIA n = 13, KO ctrl n = 14, KO MIA n = 9. Data are presented as mean ± SEM. **D, E.** Autistic score: analyzed core symptoms of ASD in females **(D)** and males **(E)**. **(D)** Two-way ANOVA genotype: F (1, 43) = 41.96; p < 0.0001, procedure: F (1, 43) = 15.54; p = 0.0003), and interaction: F (1, 43) = 10.8; p = 0.0020, followed by Tukey’s post hoc test. **(E)** Two-way ANOVA: genotype: F (1, 41) = 47.73; p < 0.0001, procedure: F (1, 41) = 8.257; p = 0.0064, and interaction: F (1, 41) = 24.02; p < 0.0001, followed by Tukey’s post hoc test. Females: WT ctrl n = 13, WT MIA n = 13, KO ctrl n = 9, KO MIA n = 12; males: WT ctrl n = 12, WT MIA n = 12, KO ctrl n = 13, KO MIA n = 8. Data are presented as mean ± SEM.

To capture the severity of the core traits associated with autism, specifically, reduced social interaction, a lower tendency for social stimuli, and increased repetitive behavior, we developed a composite behavioral score (from (Hilal et al., 2024) with modification). This method demonstrated a significant increase in autism-related behaviors in WT mice subjected to MIA and in saline-treated Lcn2 KO mice of both sexes, compared to WT control offspring (Fig. 3D, E). Notably, the combination of *Lcn2* gene deletion and prenatal immune activation did not exacerbate the behavioral phenotype. KO MIA mice exhibited similar autistic scores to both WT MIA and KO control groups (Fig. 3D, E), indicating no additive effect. This result suggests that maternal immune activation and Lcn2 deficiency may influence overlapping or converging neurodevelopmental pathways.

### Reward-motivated learning remains unchanged, and cognitive performance is unaffected following Lcn2 deletion or maternal immune activation

Cognitive impairments are a common feature of many NDDs and can be assessed through learning and memory performance (Pasciuto et al., 2015). To examine whether *Lcn2* gene deletion and maternal immune activation influence cognitive abilities, we employed a reward-motivated learning paradigm using the IntelliCage system. This automated behavioral platform allows for continuous monitoring of voluntary activity in a home-cage-like environment. In this task, mice were trained to discriminate between two drinking bottles in a designated cage corner. One bottle dispensed plain water, while the other provided a palatable sucrose solution as a positive reinforcement. The experimental design is illustrated in Fig. 4A.

**Fig. 4.**
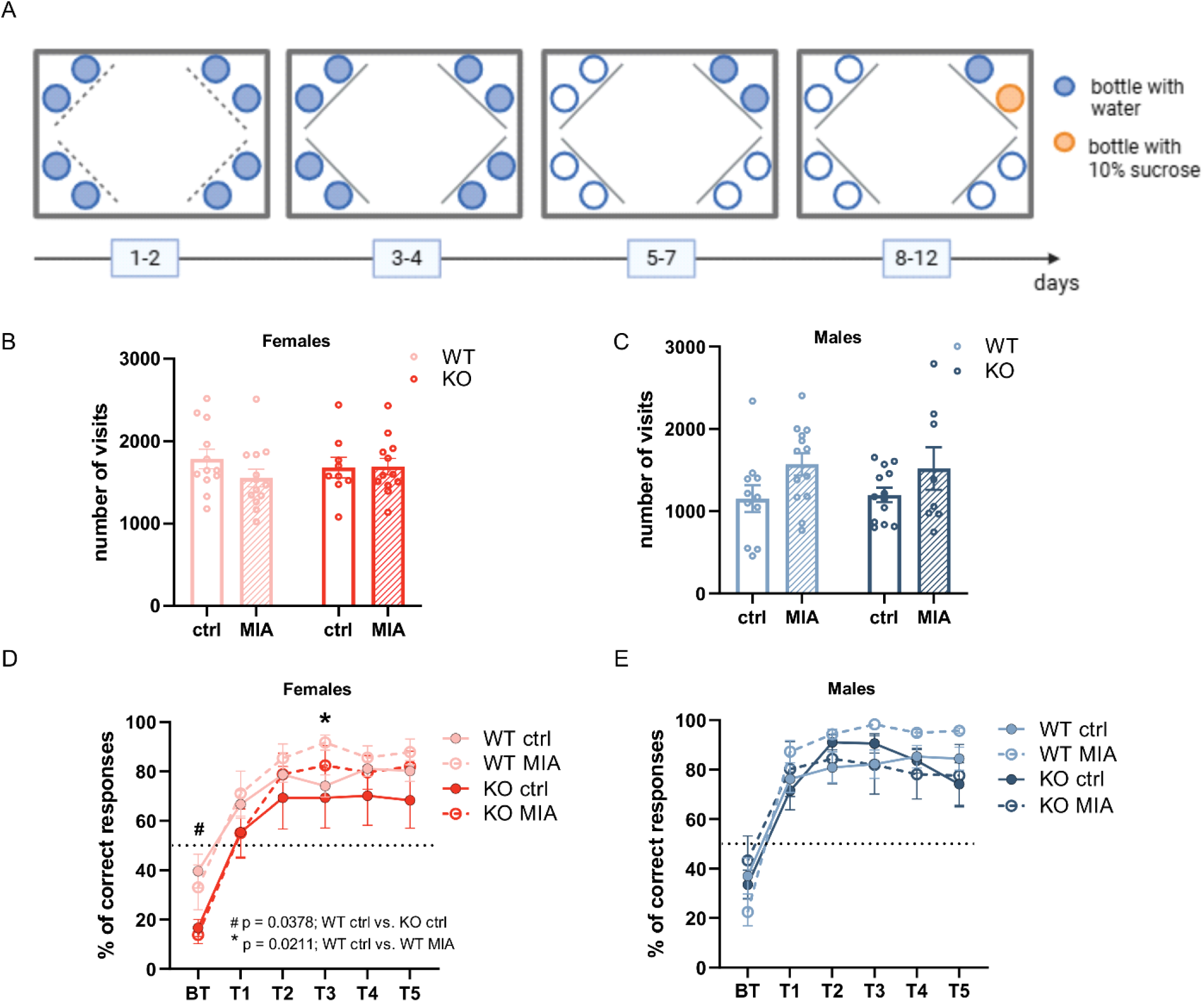
Maternal immune activation or Lcn2 deletion does not affect appetitive learning in the IntelliCage system. **A.** Timeline of the experiment in the IntelliCage. **B, C.** Activity of females **(B)** and males **(C)** presented as the number of visits to all corners during the experiment. **(B)** Two-way ANOVA: procedure: F (1, 42) = 0.9213; p = 0.3426, genotype F (1, 42) = 0.02222; p = 0.8822; and interaction: F (1, 42) = 1.136; p = 0.2925; **(C)** Two-way ANOVA: procedure: F (1, 41) = 5.680; p = 0.0219, genotype: F (1, 41) = 0.0001650; p = 0.9898; and interaction: F (1, 41) = 0.1022; p = 0.7508, followed by Tukey’s post hoc test. **D, E** Appetitive learning in females **(D)** and males **(E)** is shown as the percentage of correct responses, defined as visits in which the first nose poke was directed at the door leading to the bottle containing the sugar solution, relative to the total number of visits to that corner in a given session. **(D)** A three-way ANOVA: session: F (2.448, 102.8) = 89.93; p < 0.0001, procedure: F (1, 42) = 1.132, p = 0.2935), genotype: F (1, 42) = 3.444, p = 0.0705, interaction session x procedure: F (5, 210) = 2.388, p = 0,0392, session x genotype: F (5, 210) = 1.417, p = 0.2193, procedure x genotype: F (1, 42) = 0.01461, p = 0.9044, and session x procedure x genotype: F (5, 210) = 0.2746; p = 0.9267 with Tukey’s post hoc. **(E)** A three-way ANOVA: session: F (2,567, 105,2) = 88.58; p < 0.0001, procedure: F (1, 41) = 0.6988; p = 0.4080, genotype F (1, 41) = 0.7310; p = 0.3975, session x procedure: F (5, 205) = 0.9221; p = 0.4676, session x genotype: F (5, 205) = 3.163; p = 0.0090, procedure x genotype: F (1, 41) = 0.6314; p = 0.4314, and session x procedure x genotype: F (5, 205) = 3.925, p = 0.0020 followed by Tukey’s post hoc. PT - visits from the three days preceding the appetitive training, when water bottles were present in the corners; T1-T5 - visits from consecutive days of the training sessions. Females - WT ctrl n = 12, WT MIA n = 13, KO ctrl n = 9, KO MIA n =12; males - WT ctrl n = 11, WT MIA n = 13, KO ctrl n = 13, KO MIA n = 8. Data are presented as mean ± SEM.

Throughout the experiment, we evaluated several behavioral parameters, including general activity, learning of bottle discrimination, sucrose preference, and reward-seeking motivation (Fig. 4 and Supplementary Fig. 3). We assessed animal activity by quantifying the total number of corner visits across all experimental phases. No significant differences in overall activity were noted between experimental groups for either female or male mice (Fig. 4B, C).

Appetitive learning was assessed by calculating the percentage of visits where the first nose poke was directed at the door leading to the sucrose-containing bottle (Knapska et al., 2013). This measure was analyzed across consecutive training sessions to evaluate the acquisition of a preference for the sweetened reward. Data from the three days preceding the appetitive training (before training, BT; averaged) and the five subsequent training sessions (T1-T5) are shown (Fig. 4D, E). All experimental groups of female and male mice exhibited a significant increase in the percentage of correct responses from the before-training phase to the final training session, indicating successful learning of the location of the sucrose-containing bottle (females: WT ctrl: BT vs. T5, p = 0.001; WT MIA: p = 0.0005; KO ctrl: p = 0.0056; KO MIA: p < 0.0001; males: WT ctrl - BT vs. T5: p = 0.0016, WT MIA - BT vs. T5: p < 0.0001, KO ctrl - BT vs. T5: p = 0.0334, KO MIA - BT vs. T5: p = 0.0430, significance levels are not shown in the figure). Neither the MIA nor the *Lcn2* gene deletion affected appetitively motivated learning in either males or females (Fig. 4D, E), except on training day T3 (T3 - WT ctrl vs. WT MIA: p = 0.0211; Fig. 4D). Additionally, KO females in the control group were less likely to initiate nose pokes toward the bottle that was later replaced with the sucrose solution (BT - WT ctrl vs. KO ctrl: p = 0.0378).

To confirm that a sucrose solution served as an appetitive stimulus for the tested mice, we assessed sweet water preference, calculated as the percentage of sucrose solution consumed, measured by lick count, relative to total fluid intake during each session. In females, no significant differences in sucrose consumption between experimental groups on any training day were found (Supplementary Fig. 3A). However, all female groups demonstrated a significant increase in sucrose intake compared to the before training phase, when only plain water was available (WT ctrl - BT vs. T5: p = 0.0003; WT MIA - BT vs. T5: p = 0.0009; KO ctrl - BT vs. T5: p = 0.0005; KO MIA - BT vs. T5: p < 0.0001; significance levels not marked in the figure).

Similarly, in males, no significant differences between groups on most training days, except day 4, when KO MIA males consumed less sucrose solution than their WT MIA counterparts (T4 - WT MIA vs. KO MIA: p = 0.0435; Supplementary Fig. 3B). As observed in females, all-male groups exhibited significantly higher intake from the sucrose-containing bottle compared to the before training period with plain water (WT ctrl - BT vs. T5: p < 0.0001; WT MIA - BT vs. T5: p < 0.0001; KO ctrl - BT vs. T5: p = 0.049; KO MIA - BT vs. T5: p = 0.0379; significance levels not shown in the figure). Together, these results confirm that the sucrose solution was an appetitive stimulus for mice, as both sexes from all experimental groups consistently preferred the sweetened water.

### Lcn2 deficiencies drive transcriptomic changes in neuronal gene pathways similar to MIA

To explore the potential shared molecular mechanisms through which maternal immune activation and Lcn2 knockout affect brain function, we analyzed gene expression changes in the fetal forebrain following maternal LPS treatment. We performed bulk RNA sequencing (RNA-seq) four hours after the last LPS injection, using forebrain tissue from twelve fetuses on gestational day E18, including six controls (three males, three females) and six from LPS-treated dams (three males, three females). We focused our analysis on developing fetal forebrain. Samples from both sexes were included in the study, as no major sex-specific differences were detected in MIA- or Lcn2 knockout-associated behavioral phenotypes. To define statistically significant changes in gene expression between MIA and control offspring, as well as WT and KO animals, a false discovery rate (FDR) threshold of <0.05 was applied. Using the |log₂ fold change criteria| >1 and FDR <0.05, we observed widespread transcriptional alterations in Lcn2 KO animals (Fig. 5B), with 573 genes downregulated and 449 upregulated. In the forebrains of offspring from LPS-treated dams, 46 genes were downregulated, and 31 were upregulated (Fig. 5C). Among these, *Lcn2* mRNA showed a 2.29-fold increase (p = 0.0356); however, this change did not remain statistically significant after correction for multiple comparisons. A complete list of upregulated and downregulated gene sets is provided in Supplementary Table 1. The upregulation of *Lcn2* mRNA following MIA was subsequently confirmed by RT-qPCR (Supplementary Fig. 4).

**Fig. 5.**
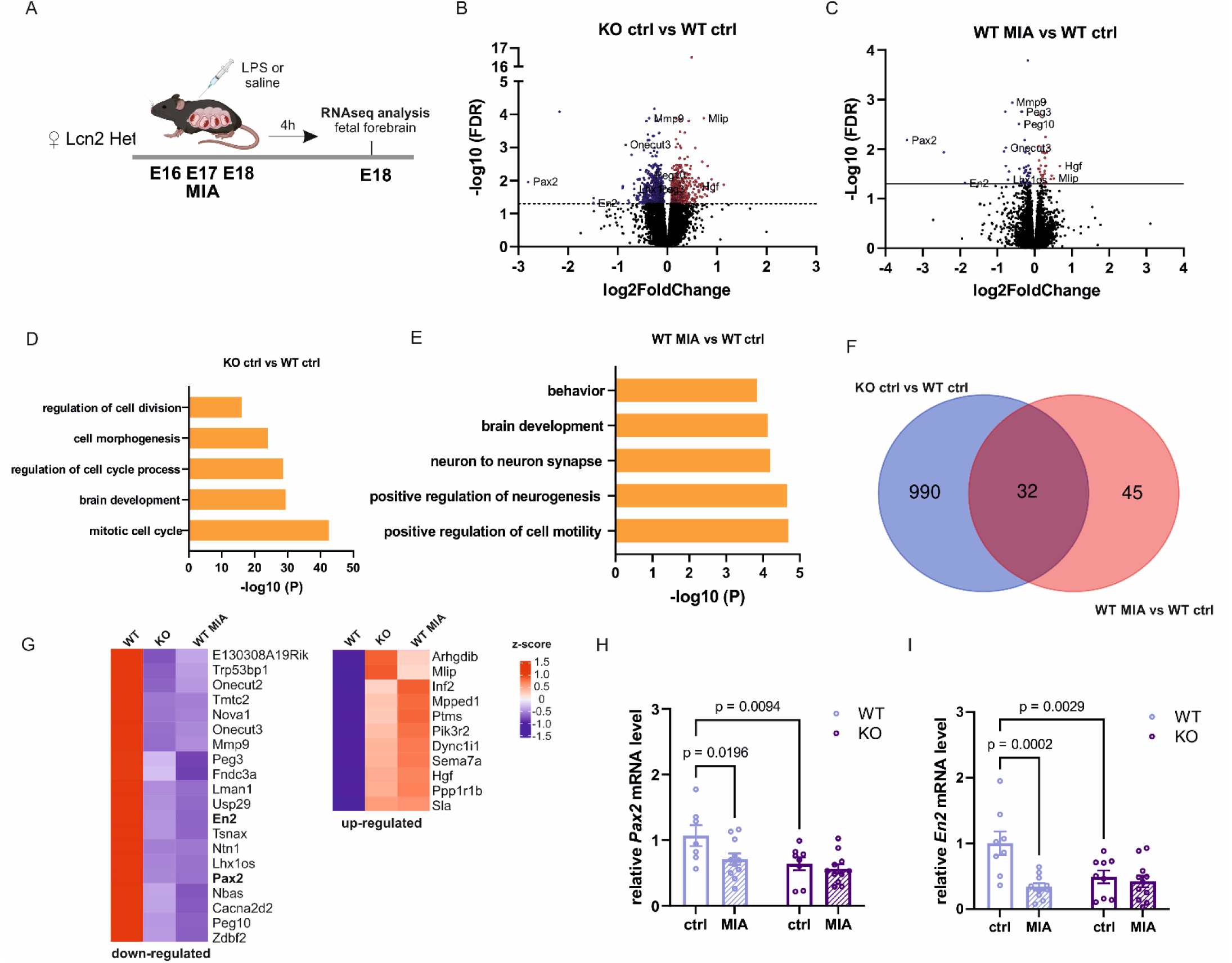
Convergent Transcriptional Signatures in Lcn2-Knockout and MIA Model. **A.** Experimental timeline. Pregnant heterozygous mice were injected with LPS or saline at E16, E17, and E18. 4 hours after the last injection, the fetal forebrain was dissected for RNA-seq analysis. **B, C.** A volcano plot illustrating DEGs in the forebrain of E18. The red dots represent significantly upregulated genes, the blue dots represent significantly downregulated genes (|log2 FC| ≥ 1 and FDR < 0.05), and the black dots represent insignificant differentially expressed genes. **(B)** DEGs in the forebrain of Lcn2 KO animals at E18 **(C)** DEGs in the forebrain of WT MIA at E18, 4 hours after the last LPS injection. **D, E.** Gene ontology enrichment analysis using Metascape based on the genes deregulated (both up- and downregulated). A bar graph showing the top 5 clusters with the highest p-value. **E.** Heatmap of enriched terms across input differentially expressed gene lists. **F.** A Venn diagram of unique and overlapping gene expression between Lcn2 KO and MIA groups. **G.** The heat map of overlapping genes between Lcn2 KO and MIA. **H, I.** RT-qPCR confirmation of the results obtained with RNAseq analysis for *Pax2* **(H)** and *En2* **(I)** genes. mRNA expression levels in the forebrain were measured 4 hours after the final injection of LPS or saline in pregnant dams. **(H)** Two-way ANOVA: procedure: F (1, 33) = 4.616; p = 0.0391, genotype: F (1, 33) = 8.232; p = 0.0071, and interaction: F (1, 33) = 1.933; p = 0.1737, followed by Fisher’s LSD test. WT ctrl n = 7, WT MIA n = 11; Lcn2 KO n = 8, Lcn2 KO MIA n = 11. **(I)** Two-way ANOVA: procedure: F (1, 34) = 11.60; p = 0.0017, genotype: F (1, 34) = 4.019; p = 0.0530, and interaction: F (1, 34) = 7.669, p = 0.0090, followed by Fisher’s LSD post hoc test. WT ctrl n = 8, WT MIA n = 10; Lcn2 KO n = 9, Lcn2 KO MIA n = 11. Data are presented as mean ± SEM.

The RNA-seq data were analyzed to identify differentially expressed genes (DEGs) between the groups and to assess the enrichment of Gene Ontology (GO) categories among these DEGs (Metascape). This analysis revealed significant enrichment in several GO categories, in DEGs in Lcn2 KO animals, and showed enrichment of ontologies associated with *brain development* (GO:0007420, 95/745 genes). In the DEGs in the MIA group, significant enrichment was also found in GO Biological Processes *brain development* (GO:0007420, 9/745genes), *behavior* (GO:0007610, 8/634), and GO Cellular Components *neuron to neuron synapse* (GO:0098984, 7/414 (Supplementary Tables 2 and 3).

Next, we examined whether the transcriptomic changes exhibited shared expression patterns between MIA and Lcn2 KO animals. Although MIA induced relatively modest changes in gene expression, we found that 40% of the differentially expressed genes in the MIA group (31 out of 77) also showed altered expression in Lcn2 KO animals (Fig. 5G). Thirty-one shared genes represented a 6.9-fold enrichment over random expectation (representation factor = 6.9, p < 7.41 × 10⁻^19^; *p* values of hypergeometric probability significance test). Functional annotation of the overlapping genes was performed through manual analysis of published literature (PubMed search), focusing on neuronal relevance. Several genes within this set are known to be involved in brain development and have been implicated in neurodevelopmental disorders, suggesting that Lcn2 deficiency and MIA may converge on common molecular pathways essential for normal brain development (Table 1). To validate the RNA-seq findings, quantitative RT-PCR was performed for selected genes associated with an increased risk for autism spectrum disorder (Fig. 5H, I). MIA resulted in a significant downregulation of *Pax2* and *En2* mRNA expression in the forebrain of fetuses as compared to controls (Fig. 5H, I) (*Pax2*: WT ctrl vs. WT MIA p = 0.0196, *En2* WT ctrl vs. WT MIA p = 0.0002). Similar downregulation of *Pax2* and *En2* mRNA was observed in Lcn2 KO animals (*Pax2*: WT ctrl vs. KO ctrl, p = 0.0094, *En2* WT ctrl vs. KO ctrl, p = 0.0029).

**Table 1.** DEGs regulating brain development with a link to neurodevelopmental disorders in humans.

| **GENE** | **Name** | **Function** | **Link with neurodevelopmental disorders** | **References** |
| --- | --- | --- | --- | --- |
| **Pax2** | Paired Box 2 | transcription factor, present in the developing brain, regulates the differentiation of GABAergic neurons | autism spectrum disorder, intellectual disability, epilepsy | (Lv et al., 2021; Rossanti et al., 2020) |
| **EN2** | engrailed homeobox 2 | transcription factor, critical for brain development, regulates the number of GABAergic neurons | autism spectrum disorder | (Benayed et al., 2009; Benayed et al., 2005; Gharani et al., 2004; Provenzano et al., 2020; Sen et al., 2010; Sgado et al., 2013) |
| **Hgf** | Hepatocyte Growth Factor | growth factor, neurotrophic factor involved in synaptogenesis | autism spectrum disorder | (Desole et al., 2021; Galvez-Contreras et al., 2017; Sugihara et al., 2007) |
| **Mmp9** | Matrix metalloproteinase 9 | metalloproteinase, regulates functional and structural plasticity | autism spectrum disorder, fragile X syndrome, schizophrenia | (Dziembowska et al., 2013; Legutko et al., 2024; Lepeta et al., 2017; Michaluk et al., 2011; Szepesi et al., 2013) |
| **Nova1** | alternative splicing regulator 1 | RNA-binding protein, neuronal splicing factor, controls transcript localization and regulation of transcript stability | autism | (Tajima et al., 2023; Zhang et al., 2010) |
| **Sema7a** | Semaphorin 7A | is a key regulator of axon guidance, synapse formation | autism, developmental delay | (Carcea et al., 2014; Jongbloets et al., 2017) |
| **Ppp1r1b**<br>(DARPP-32) | protein phosphatase 1 regulatory inhibitor subunit 1B | inhibitor of protein-phosphatase 1, plays a key role in neuronal signaling, particularly within dopamine pathways, regulates the number of GABAergic neurons | autism, schizophrenia | (Akkouh et al., 2024; Hettinger et al., 2012; Souza et al., 2024) |
| **Cacna2d2** | alpha-2/delta-2 ($\alpha 2\delta 2$) subunit of voltage-gated calcium channels | voltage-gated calcium channel, controls calcium-dependent signaling in neurons | epilepsy, schizophrenia | (Ablinger et al., 2020; Punetha et al., 2019) |
| **Ntn1** | netrin-1 | secreted protein, cue for axon guidance, promotes axonal branching, regulates the number and function of excitatory synapses | no direct link | (Goldman et al., 2013; Nabeel Mustafa et al., 2025; Serafini et al., 1996) |

## Discussion

This study suggests that Lcn2 is a regulator of brain development. We demonstrated that Lcn2 is expressed in the brain during development and is upregulated following MIA, particularly in the hippocampus and neocortex. Behavioral analyses have revealed that Lcn2 deletion and MIA independently disrupted social behaviors and increased repetitive behaviors in offspring without additive or synergistic effects, suggesting a potential occlusion of shared pathways. Notably, these behavioral alterations were selective, as cognitive performance in learning and memory tasks remained unaffected. Transcriptomic analysis of the fetal forebrain further supports this convergence, revealing a shared set of differentially expressed genes in both Lcn2-knockout and MIA-exposed fetal brains. These results suggest that while increased Lcn2 expression following prenatal immune challenge does not appear to be the primary driver of behavioral disruptions in this model, Lcn2 likely plays an essential role in supporting neurodevelopment under physiological conditions.

While Lcn2 has been extensively studied in the context of adult brain pathologies, such as neuroinflammation and neurodegenerative processes (Doliwa et al., 2024; Ferreira et al., 2015; Ip et al., 2011; Kuzniewska et al., 2016; Mucha et al., 2011), its role during fetal brain development has remained largely unexplored. Our study uncovers a novel role for Lcn2 in shaping brain development. We demonstrate that Lcn2 expression in the mouse hippocampus is developmentally regulated, with mRNA levels detectable prenatally and peaking perinatally. Lcn2 expression has also been observed in the developing human brain (Zhang et al., 2012) and in zebrafish embryos (Lee et al., 2009), indicating a potentially conserved role for this protein in neurodevelopment across species. Although no direct evidence for the role of Lcn2 in human neurodevelopmental disorders has been provided so far, it is important to note that either deletion or insertion identified within the 486.52–703.2 kb genomic region that encompasses the *Lcn2* gene in humans (according to the DECIPHER database; https://www.deciphergenomics.org/) has been found in patients with neurodevelopmental disorders’ symptoms including autism, global developmental delay, and intellectual disability.

Previous studies have shown that *Lcn2* expression increases in the adult mouse brain following systemic LPS administration (Hamzic et al., 2013; Kang et al., 2018; Marques et al., 2008). In the current study, we demonstrate for the first time that MIA induced by repeated low-dose LPS injections during late gestation leads to a robust upregulation of *Lcn2* mRNA and protein in the fetal neocortex and hippocampus. This increase is accompanied by elevated maternal serum Lcn2 levels, indicating a systemic inflammatory response associated with MIA. A recent study also reported elevated Lcn2 levels in both the blood and hippocampus of mice at postnatal day 4, 24 hours after bacterial infection with *Staphylococcus epidermidis* (Gravina et al., 2023).

Social behavior deficits are among the most consistently observed phenotypes in maternal immune activation models (Choi et al., 2016; Dutra et al., 2023; Fernandez de Cossio et al., 2017; Kentner et al., 2019; Malkova et al., 2012; Mattei et al., 2014; Wu et al., 2018). To assess the impact of MIA and *Lcn2* gene deletion on social behavior in mice, we used the Eco-HAB® system, a validated tool for detecting social deficits in rodents (Puścian et al., 2016; Puścian et al., 2022; Roszkowska et al., 2022; Szewczyk et al., 2024). It enables continuous tracking of spontaneous, long-term social interactions among group-housed animals in a seminaturalistic environment, providing an ethologically relevant measure of sociability. Using this system, we have found that MIA disrupts sociability in wild-type offspring, as male and female mice born to LPS-treated dams were less willing to spend time with conspecifics than control mice. Interestingly, deletion of the *Lcn2* gene alone in control mice of both sexes produced a similar decrease in in-cohort sociability. Notably, Lcn2 deletion alone had a greater impact on social behavior in females than prenatal LPS exposure. Similar deficits in in-cohort sociability have been reported in various animal models of autism spectrum disorder, further supporting the relevance of the findings (Puścian et al., 2016; Rydzanicz et al., 2024).

Reduced interest in social stimuli, another marker of social dysfunction, was analyzed in the Eco-HAB**®** and modified three-chamber tests (Szewczyk et al., 2024). In both tests, MIA decreased social interest in WT females, while Lcn2 deletion reduced social preference only in males. In addition, the three-chamber test revealed an effect of LPS-induced reduction in interest toward the social stimulus in Lcn2 KO in females, which was not observed in Eco-HAB**®**. These discrepancies are probably due to differences in testing conditions, such as animals being assessed individually versus in a group, or variations in test duration. Nevertheless, these findings highlight that Lcn2 deficiency alone can impair preference for social stimuli, mimicking MIA-induced phenotypes. Similar reductions in social interest have been reported in offspring across various MIA models (Kentner et al., 2019).

Besides impaired social interactions, repetitive behaviors are among the most consistently reported phenotypes following maternal immune activation (Choi et al., 2016; Coiro et al., 2015; Fernandez de Cossio et al., 2017; Malkova et al., 2012; Pendyala et al., 2017). Using the marble burying test, a well-established measure of repetitive behavior, we found that *Lcn2* knockout increased the number of marbles buried in both sexes. Interestingly, prenatal LPS exposure increased marble burying only in wild-type males, indicating a sex- and genotype-dependent effect. However, in Lcn2 KO males and females, MIA did not further alter this behavior. The absence of additive effects in KO mice exposed to MIA implies that Lcn2-dependent pathways may be a critical point of convergence in MIA-induced behavioral alterations. A similar phenomenon has been reported in Fmr1 KO mice, where Poly(I:C)-induced MIA did not exacerbate core autism-like behaviors (Hilal et al., 2024).

Previous studies have reported inconclusive findings regarding the impact of MIA on cognitive function in offspring, with some reporting learning impairments following prenatal immune challenge, while others observed no significant deficits in learning and memory (Abazyan et al., 2010; Batinić et al., 2016; Giovanoli et al., 2015; Golan et al., 2005). In our study, no significant effect of either MIA or *Lcn2* gene deletion on appetitively motivated spatial learning was found. Although learning deficits have previously been reported in Lcn2 knockout mice (Ferreira et al., 2013; Ferreira et al., 2018; Ferreira et al., 2023), our Lcn2 knockout controls were also exposed to prenatal injection stress. This factor may have impacted brain development and contributed to phenotypic differences from previous studies. These findings suggest that Lcn2 and MIA may preferentially influence neural circuits underlying social and repetitive behaviors rather than broadly affect cognitive function.

The mechanisms by which prenatal infection disrupts neurodevelopment are still not fully understood. In this study, we compared gene expression profiles between MIA-exposed animals and Lcn2 knockout mice, which exhibit similar behavioral impairments. In both experimental groups, among differentially expressed genes (DEGs), gene ontology analysis showed enrichment in genes in the *brain development* category. These findings are in line with previous MIA transcriptomic studies identifying genes in specific categories related to central nervous system development (Canales et al., 2021; Lombardo et al., 2018; Oskvig et al., 2012). We found a 6.9-fold enrichment of overlap genes between MIA and Lcn2 KO DEGs. On the molecular level, functional annotation of the shared genes indicated their role in control of inhibitory/excitatory balance and the regulation of functional and structural plasticity, highlighting their potential involvement in shaping neural circuit function (see Table 1). Our results are consistent with other studies showing that MIA may impact the GABAergic transmission by decreasing the number of GABAergic neurons (Nouel et al., 2012; Shi et al., 2009; Wischhof et al., 2015; Zhang and van Praag, 2015). Notably, animal models have also demonstrated that offspring exposed to maternal immune challenges exhibit synaptic dysfunction and reduced synaptic plasticity (Hui et al., 2020; Ito et al., 2010). Moreover, mutations in many of the genes identified in our study are associated with increased susceptibility to autism spectrum disorder and other neurodevelopmental disorders in humans (Table 1).

Despite the identification of shared transcriptional programs across our experimental groups, the signaling pathways underlying the convergence of these transcriptional changes remain unclear. Remarkably, even without an external inflammatory stimulus, Lcn2 deficiency may prime the fetal brain toward a transcriptional state characteristic of prenatal immune activation. However, the role of Lcn2 as an immunomodulatory protein remains controversial. While some studies have identified Lcn2 as a proinflammatory mediator that amplifies neuroinflammation (Behrens et al., 2021; Jin et al., 2014; Kim et al., 2024; Lee et al., 2011; Lee et al., 2009; Li et al., 2023; Liu et al., 2022; Shin et al., 2021), other findings suggest a more complex role. For example, lack of Lcn2 has been associated with increased TNFα and IL-6 expression and enhanced LPS-induced behaviors, suggesting a potential anti-inflammatory role (Kang et al., 2018). However, other studies report no significant differences in brain inflammation between wild-type and Lcn2 knockout mice following LPS exposure (Gasterich et al., 2021; Vichaya et al., 2019). Interestingly, both the absence of Lcn2 and its elevated levels appear to produce similar outcomes, possibly depending on the specific context in which Lcn2 is upregulated.

While our findings provide novel molecular insights into the effects of MIA and Lcn2 on fetal brain development, they should be interpreted considering the limitations of the methods. Although bulk RNA sequencing offered valuable insights into shared molecular changes, single-cell transcriptomics and spatial transcriptomics in fetal forebrain tissues of Lcn2 KO and MIA mice would clarify the cellular origin of DEGs. In particular, the origin of Lcn2 expression during development and under MIA conditions remains unresolved. In physiological and pathological conditions, Lcn2 has been detected in multiple cell types, including astrocytes, microglia, neurons, epithelial cells, and choroid plexus endothelial cells (Chia et al., 2011; Doliwa et al., 2024; Ip et al., 2011; Kim et al., 2024; Mucha et al., 2011). However, which of these cell types contributes to Lcn2 expression in the prenatal brain during neuroinflammatory events is unknown. Another potential limitation of our study is the relatively modest number of genes differentially expressed following MIA. Although repeated LPS injections model MIA during key brain developmental stages, including neurogenesis and synaptogenesis (Khalaf-Nazzal and Francis, 2013; Li et al., 2010), they may also induce immune tolerance, dampening inflammatory and transcriptional responses (Chen et al., 2005; Fan and Cook, 2004). This phenomenon could partially explain the limited number of differentially expressed genes detected in the MIA condition.

In conclusion, we have demonstrated that Lcn2 is a key factor in brain development, particularly in the formation of circuits that underlie social and repetitive behavior. The similar molecular and behavioral outcomes observed with Lcn2 deletion and MIA indicate that these interactions may converge on common developmental pathways.

## Acknowledgments

This work was supported by the National Science Centre, Poland: NCN grant 2017/27/B/NZ4/01639 and 2022/45/B/NZ4/03262. The authors report neither financial interests nor conflicts of interest.

## Author contributions

**Martyna Pekala**: conceptualization, investigation; methodology, visualization, writing original draft; **Sylwia Zawiślak**: investigation, visualization; **Sandra Romanis**: investigation, visualization; **Karolina Nader**: investigation, visualization; **Aleksandra Cabaj**: investigation, methodology; **Anna Madecka**: methodology; **Alicja Puścian**: methodology; **Ewelina Knapska**: methodology; **Robert Pawlak:** resources; **Leszek Kaczmerek**: resources; **Katarzyna Kalita:** conceptualization, funding acquisition, methodology, supervision, project administration, writing original draft.

---

**Supplementary Fig. 1.**
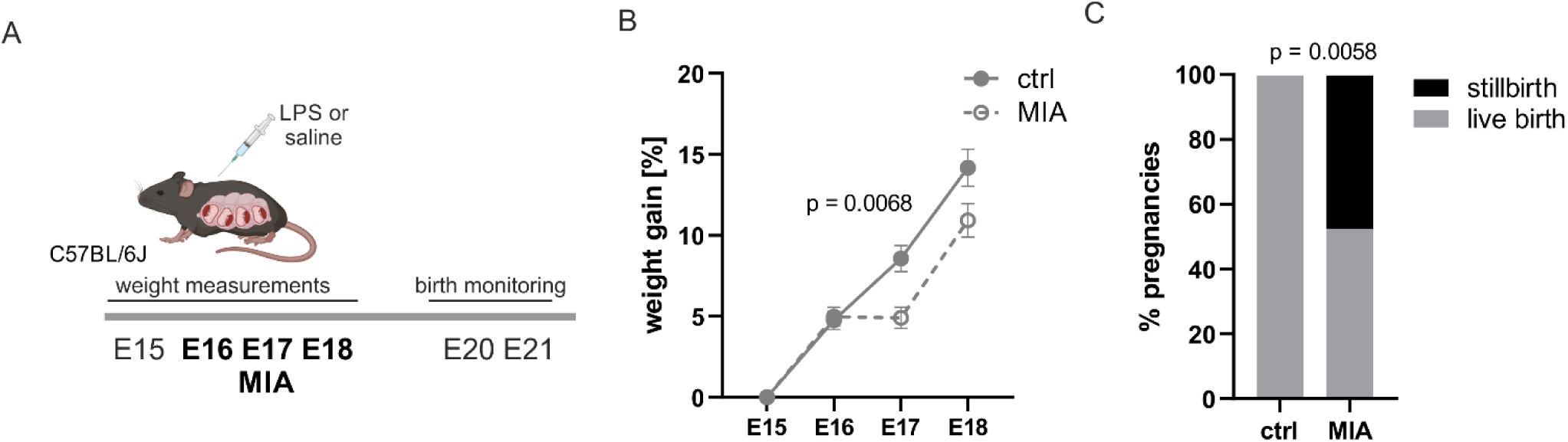
MIA affects weight gain in pregnant C57BL/6J females and fetal survival. **A.** Scheme of the experiment. LPS was administered to pregnant C57BL/6J mice on embryonic days E16, E17, and E18. Pregnant females’ weight gain and offspring survival were analysed. **B.** Percentage of weight gain in pregnant females relative to their body weight on the day before the first injection. WT ctrl n = 11, WT MIA n = 12. Two-way repeated measures ANOVA: treatment: F (1, 20) = 4.939; p = 0.0379, time: F (1.437, 28.75 = 114.6; p < 0.0001, and interaction: F (2, 40) = 8.147; p = 0.0011, followed by Sidak’s post hoc test. Data are presented as mean ± SEM. **C.** Fetal survival, shown as the percentage of pregnancies resulting in live or stillbirths. WT ctrl n = 11, WT MIA n = 21. χ² test analysis.

**Supplementary Fig. 2.**
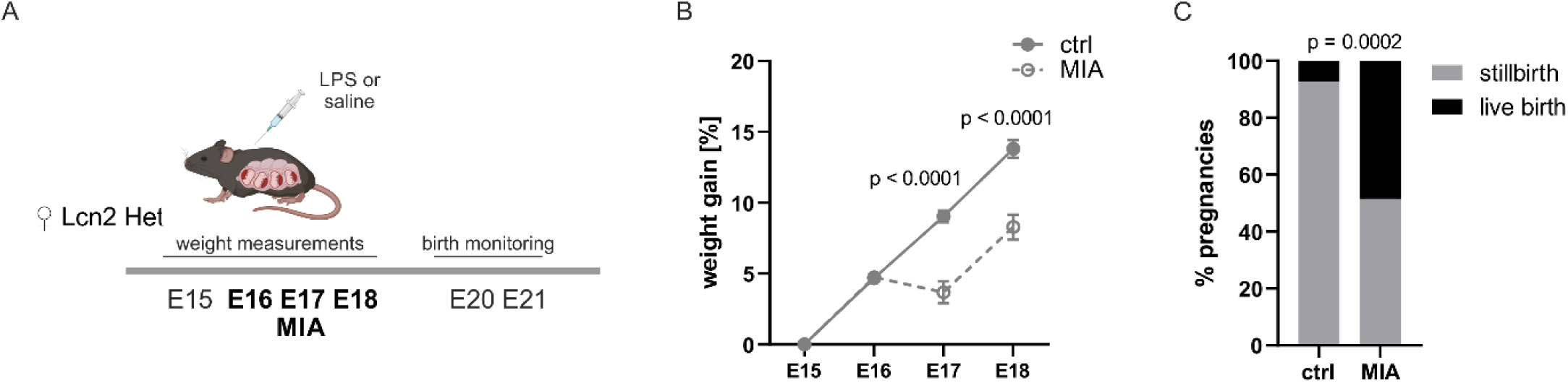
Effect of MIA on weight gain in pregnant Lcn2 Het females and fetal survival. **A.** Scheme of the experiment. LPS was administered to pregnant Lcn2 Het mice on embryonic days E16, E17, and E18. Pregnant females’ weight gain and offspring survival were analyzed. **B.** Percentage of weight gain in pregnant females relative to their body weight on the day before the first injection. Ctrl n = 28, MIA = 34. Two-way repeated measures ANOVA: treatment: F (1, 60) = 26.89; p < 0.0001, time: F (1.633, 97.97) = 104.3; p < 0,0001, and interaction: F (2, 120) = 24.39; p < 0.0001, followed by Sidak’s post hoc test. Data are presented as mean ± SEM. **C.** Fetal survival is shown as the percentage of Lcn2 KO pregnancies resulting in live or stillbirths. WT ctrl n = 27, WT MIA n = 66. χ² test analysis.

**Supplementary Fig. 3.**
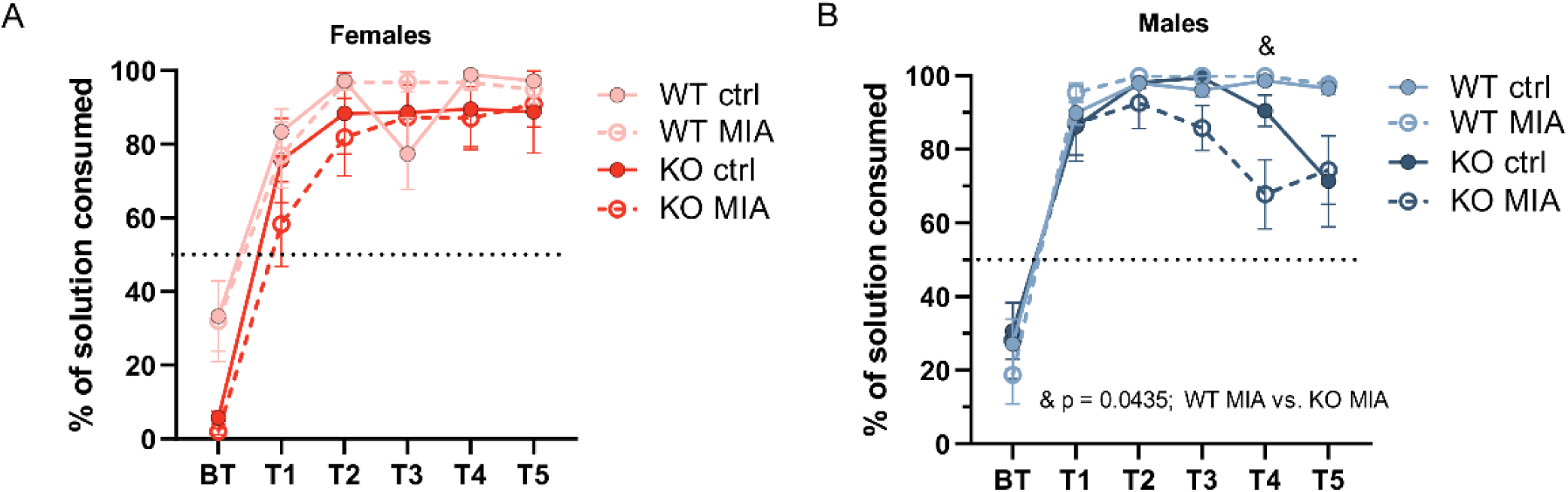
Maternal immune activation or Lcn2 deletion does not influence sucrose consumption. Sucrose consumption in females **(A)** and males **(B)** in IntelliCage. Preference for sweetened as the percentage of sugar solution consumed (measured by the number of licks) relative to the total fluid intake during a given session. **(A)** A three-way ANOVA: session F (2.198, 92.32) = 104.2; p < 0.0001, procedure F (1, 42) = 0.1118; p = 0.7397, genotype F (1, 42) = 4.195; p = 0.0468, session x procedure F (5, 210) = 1.378; p = 0.2337, session x genotype F (5, 210) = 2.955; p = 0.0134, procedure x genotype F (1, 42) = 0.2878; p = 0.5945, session x procedure x genotype F (5, 210) = 0.6052; p = 0.6960, followed by Tukey’s post hoc test. Data are presented as mean ± SEM. **(B)** A three-way ANOVA: session: F (2.297, 94.18) = 104.0; p < 0.0001, procedure: F (1, 41) = 0.8240; p = 0.3693; genotype: F (1, 41) = 7.366; p = 0.0097, session x procedure F (5, 205) = 0.9636; p = 0.4412, session x genotype F (5, 205) = 4.624; p = 0.0005, procedure x genotype F (1, 41) = 1.377; p = 0.2474, session x procedure x genotype F (5, 205) = 1.163; p = 0.3288, followed by Tukey’s post hoc test. Data are presented as mean ± SEM.

**Supplementary Fig. 4.**
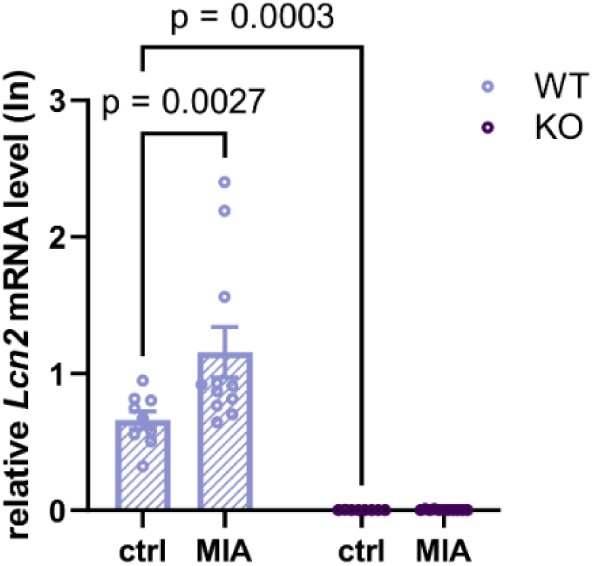
Lcn2 expression in the forebrain of WT and Lcn2 KO animals at h after MIA. *Lcn2* mRNA expression levels in fetal forebrain, measured 4 hours after the final injection of LPS or saline in pregnant dams. Data were log-transformed (Y = Ln(Y)) before analysis. WT ctrl n = 9, WT MIA n = 11; Lcn2 KO n = 8, Lcn2 KO MIA n = 11. Two-way ANOVA: procedure: F (1, 35) = 5.079; p = 0.0306, genotype: F (1, 35) = 67.80; p < 0.0001, interaction: F (1, 35) = 5.023; p = 0.0314, Fisher’s LSD post hoc test. Data are presented as mean ± SEM.

## References

1. Abazyan, B., Nomura, J., Kannan, G., Ishizuka, K., Tamashiro, K.L.K., Nucifora, F., Pogorelov, V., Ladenheim, B., Yang, C., Krasnova, I.N., et al. (2010). Prenatal interaction of mutant DISC1 and immune activation produces adult psychopathology. Biol. Psychiatry 68, 1172–1181.

2. Ablinger, C., Geisler, S.M., Stanika, R.I., Klein, C.T., and Obermair, G.J. (2020). Neuronal alpha(2)delta proteins and brain disorders. Pflugers Arch 472, 845–863.

3. Akkouh, I.A., Ueland, T., Szabo, A., Hughes, T., Smeland, O.B., Andreassen, O.A., Osete, J.R., and Djurovic, S. (2024). Longitudinal Transcriptomic Analysis of Human Cortical Spheroids Identifies Axonal Dysregulation in the Prenatal Brain as a Mediator of Genetic Risk for Schizophrenia. Biol Psychiatry 95, 687–698.

4. Ayubi, E., and Mansori, K. (2022). Maternal Infection during Pregnancy and Attention-Deficit Hyperactivity Disorder in Children: A Systematic Review and Meta-Analysis. Iran J Public Health 51, 2674–2687.

5. Batinić, B., Santrač, A., Divović, B., Timić, T., Stanković, T., Obradović, A.L., Joksimović, S., and Savić, M.M. (2016). Lipopolysaccharide exposure during late embryogenesis results in diminished locomotor activity and amphetamine response in females and spatial cognition impairment in males in adult, but not adolescent rat offspring. Behavioural Brain Research 299, 72–80.

6. Behrens, V., Voelz, C., Müller, N., Zhao, W., Gasterich, N., Clarner, T., Beyer, C., and Zendedel, A. (2021). Lipocalin 2 as a Putative Modulator of Local Inflammatory Processes in the Spinal Cord and Component of Organ Cross talk After Spinal Cord Injury. Molecular neurobiology 58, 5907–5919.

7. Benayed, R., Choi, J., Matteson, P.G., Gharani, N., Kamdar, S., Brzustowicz, L.M., and Millonig, J.H. (2009). Autism-associated haplotype affects the regulation of the homeobox gene, ENGRAILED 2. Biol Psychiatry 66, 911–917.

8. Benayed, R., Gharani, N., Rossman, I., Mancuso, V., Lazar, G., Kamdar, S., Bruse, S.E., Tischfield, S., Smith, B.J., Zimmerman, R.A., et al. (2005). Support for the Homeobox Transcription Factor Gene ENGRAILED 2 as an Autism Spectrum Disorder Susceptibility Locus. The American Journal of Human Genetics 77, 851–868.

9. Boksa, P. (2010). Effects of prenatal infection on brain development and behavior: a review of findings from animal models. Brain Behav Immun 24, 881–897.

10. Bosch-Bouju, C., Larrieu, T., Linders, L., Manzoni, O.J., and Laye, S. (2016). Endocannabinoid-Mediated Plasticity in Nucleus Accumbens Controls Vulnerability to Anxiety after Social Defeat Stress. Cell Rep 16, 1237–1242.

11. Canales, C.P., Estes, M.L., Cichewicz, K., Angara, K., Aboubechara, J.P., Cameron, S., Prendergast, K., Su-Feher, L., Zdilar, I., Kreun, E.J., et al. (2021). Sequential perturbations to mouse corticogenesis following in utero maternal immune activation. Elife 10.

12. Carcea, I., Patil, S.B., Robison, A.J., Mesias, R., Huntsman, M.M., Froemke, R.C., Buxbaum, J.D., Huntley, G.W., and Benson, D.L. (2014). Maturation of cortical circuits requires Semaphorin 7A. Proceedings of the National Academy of Sciences of the United States of America 111, 13978–13983.

13. Chang, Y.C., Cole, T.B., and Costa, L.G. (2017). Behavioral Phenotyping for Autism Spectrum Disorders in Mice. Curr Protoc Toxicol 72, 112211–112221.

14. Chen, R., Zhou, H., Beltran, J., Malellari, L., and Chang, S.L. (2005). Differential expression of cytokines in the brain and serum during endotoxin tolerance. J. Neuroimmunol. 163, 53–72.

15. Chia, W.J., Dawe, G.S., and Ong, W.Y. (2011). Expression and localization of the iron-siderophore binding protein lipocalin 2 in the normal rat brain and after kainate-induced excitotoxicity. Neurochem Int 59, 591–599.

16. Choi, G.B., Yim, Y.S., Wong, H., Kim, S., Kim, H., Kim, S.V., Hoeffer, C.A., Littman, D.R., and Huh, J.R. (2016). The maternal interleukin-17a pathway in mice promotes autism-like phenotypes in offspring. Science (New York, N.Y.) 351, 933–939.

17. Coiro, P., Padmashri, R., Suresh, A., Spartz, E., Pendyala, G., Chou, S., Jung, Y., Meays, B., Roy, S., Gautam, N., et al. (2015). Impaired synaptic development in a maternal immune activation mouse model of neurodevelopmental disorders. Brain Behav. Immun. 50, 249–258.

18. Desole, C., Gallo, S., Vitacolonna, A., Montarolo, F., Bertolotto, A., Vivien, D., Comoglio, P., and Crepaldi, T. (2021). HGF and MET: From Brain Development to Neurological Disorders. Front Cell Dev Biol 9, 683609.

19. Doliwa, M., Kuzniewska, B., Nader, K., Reniewicz, P., Kaczmarek, L., Michaluk, P., and Kalita, K. (2024). Astrocyte-Secreted Lcn2 Modulates Dendritic Spine Morphology [v1].

20. Dunaevsky, A., and Bergdolt, L. (2019). Brain changes in a maternal Immune activation model of neurodevelopmental brain disorders. Progress in Neurobiology 175, 1–19.

21. Dutra, M.L., Dias, P., Freiberger, V., Ventura, L., Comim, C.M., Martins, D.F., and Bobinski, F. (2023). Maternal immune activation induces autism-like behavior and reduces brain-derived neurotrophic factor levels in the hippocampus and offspring cortex of C57BL/6 mice. Neuroscience Letters 793, 136974.

22. Dziembowska, M., Pretto, D.I., Janusz, A., Kaczmarek, L., Leigh, M.J., Gabriel, N., Durbin-Johnson, B., Hagerman, R.J., and Tassone, F. (2013). High MMP-9 activity levels in fragile X syndrome are lowered by minocycline. Am. J. Med. Genet. 161, 1897–1903.

23. Fan, H., and Cook, J.A. (2004). Molecular mechanisms of endotoxin tolerance. Journal of Endotoxin Research 10, 71–84.

24. Fernandez de Cossio, L., Guzman, A., van der Veldt, S., and Luheshi, G.N. (2017). Prenatal infection leads to ASD-like behavior and altered synaptic pruning in the mouse offspring. Brain Behav Immun 63, 88–98.

25. Ferreira, A.C., Dá Mesquita, S., Sousa, J.C., Correia-Neves, M., Sousa, N., Palha, J.A., and Marques, F. (2015). From the periphery to the brain: Lipocalin-2, a friend or foe? Progress in Neurobiology 131, 120–136.

26. Ferreira, A.C., Pinto, V., Mesquita, S.D., Novais, A., Sousa, J.C., Correia-Neves, M., Sousa, N., Palha, J.A., and Marques, F. (2013). Lipocalin-2 is involved in emotional behaviors and cognitive function. Front. Cell. Neurosci. 7.

27. Ferreira, A.C., Santos, T., Sampaio-Marques, B., Novais, A., Mesquita, S.D., Ludovico, P., Bernardino, L., Correia-Neves, M., Sousa, N., Palha, J.A., et al. (2018). Lipocalin-2 regulates adult neurogenesis and contextual discriminative behaviours. Molecular Psychiatry 23, 1031–1039.

28. Ferreira, A.C., Sousa, N., Sousa, J.C., and Marques, F. (2023). Age-related changes in mice behavior and the contribution of lipocalin-2. Front. Aging Neurosci. 15.

29. Flo, T.H., Smith, K.D., Sato, S., Rodriguez, D.J., Holmes, M.A., Strong, R.K., Akira, S., and Aderem, A. (2004). Lipocalin 2 mediates an innate immune response to bacterial infection by sequestrating iron. Nature 432, 917–921.

30. Galvez-Contreras, A.Y., Campos-Ordonez, T., Gonzalez-Castaneda, R.E., and Gonzalez-Perez, O. (2017). Alterations of Growth Factors in Autism and Attention-Deficit/Hyperactivity Disorder. Front Psychiatry 8, 126.

31. Gao, Y., Slomnicki, L.P., Kilanczyk, E., Forston, M.D., Pietrzak, M., Rouchka, E.C., Howard, R.M., Whittemore, S.R., and Hetman, M. (2024). Reduced Expression of Oligodendrocyte Linage-Enriched Transcripts During the Endoplasmic Reticulum Stress/Integrated Stress Response. ASN Neuro 16, 2371162.

32. Gasterich, N., Wetz, S., Tillmann, S., Fein, L., Seifert, A., Slowik, A., Weiskirchen, R., Zendedel, A., Ludwig, A., Koschmieder, S., et al. (2021). Inflammatory Responses of Astrocytes Are Independent from Lipocalin 2. J Mol Neurosci 71, 933–942.

33. Gharani, N., Benayed, R., Mancuso, V., Brzustowicz, L.M., and Millonig, J.H. (2004). Association of the homeobox transcription factor, ENGRAILED 2, 3, with autism spectrum disorder. Mol Psychiatry 9, 474–484.

34. Giovanoli, S., Notter, T., Richetto, J., Labouesse, M.A., Vuillermot, S., Riva, M.A., and Meyer, U. (2015). Late prenatal immune activation causes hippocampal deficits in the absence of persistent inflammation across aging. Journal of Neuroinflammation 12.

35. Goetz, D.H., Holmes, M.A., Borregaard, N., Bluhm, M.E., Raymond, K.N., and Strong, R.K. (2002). The Neutrophil Lipocalin NGAL Is a Bacteriostatic Agent that Interferes with Siderophore-Mediated Iron Acquisition. Molecular Cell 10, 1033–1043.

36. Golan, H.M., Lev, V., Hallak, M., Sorokin, Y., and Huleihel, M. (2005). Specific neurodevelopmental damage in mice offspring following maternal inflammation during pregnancy. Neuropharmacology 48, 903–917.

37. Goldman, J.S., Ashour, M.A., Magdesian, M.H., Tritsch, N.X., Harris, S.N., Christofi, N., Chemali, R., Stern, Y.E., Thompson-Steckel, G., Gris, P., et al. (2013). Netrin-1 promotes excitatory synaptogenesis between cortical neurons by initiating synapse assembly. The Journal of neuroscience : the official journal of the Society for Neuroscience 33, 17278–17289.

38. Gravina, G., Ardalan, M., Chumak, T., Nilsson, A.K., Ek, J.C., Danielsson, H., Svedin, P., Pekny, M., Pekna, M., Sävman, K., et al. (2023). Proteomics identifies lipocalin-2 in neonatal inflammation associated with cerebrovascular alteration in mice and preterm infants. iScience 26.

39. Hamzic, N., Blomqvist, A., and Nilsberth, C. (2013). Immune-Induced Expression of Lipocalin-2 in Brain Endothelial Cells: Relationship with Interleukin-6, Cyclooxygenase-2 and the Febrile Response. Journal of Neuroendocrinology 25, 271–280.

40. Han, V.X., Patel, S., Jones, H.F., and Dale, R.C. (2021a). Maternal immune activation and neuroinflammation in human neurodevelopmental disorders. Nature Reviews Neurology 17, 564–579.

41. Han, V.X., Patel, S., Jones, H.F., Nielsen, T.C., Mohammad, S.S., Hofer, M.J., Gold, W., Brilot, F., Lain, S.J., Nassar, N., et al. (2021b). Maternal acute and chronic inflammation in pregnancy is associated with common neurodevelopmental disorders: a systematic review. Transl Psychiatry 11, 71.

42. Hettinger, J.A., Liu, X., Hudson, M.L., Lee, A., Cohen, I.L., Michaelis, R.C., Schwartz, C.E., Lewis, S.M., and Holden, J.J. (2012). DRD2 and PPP1R1B (DARPP-32) polymorphisms independently confer increased risk for autism spectrum disorders and additively predict affected status in male-only affected sib-pair families. Behav Brain Funct 8, 19.

43. Hilal, M.L., Rosina, E., Pedini, G., Restivo, L., and Bagni, C. (2024). Dysregulation of the mTOR-FMRP pathway and synaptic plasticity in an environmental model of ASD. Molecular Psychiatry, 1–15.

44. Hooley, J.M. (2010). Social Factors in Schizophrenia. Curr Dir Psychol Sci 19, 238–242.

45. Hui, C.W., Vecchiarelli, H.A., Gervais, É., Luo, X., Michaud, F., Scheefhals, L., Bisht, K., Sharma, K., Topolnik, L., and Tremblay, M.-È. (2020). Sex Differences of Microglia and Synapses in the Hippocampal Dentate Gyrus of Adult Mouse Offspring Exposed to Maternal Immune Activation. Front. Cell. Neurosci. 14.

46. Ip, J.P.K., Noçon, A.L., Hofer, M.J., Lim, S.L., Müller, M., and Campbell, I.L. (2011). Lipocalin 2 in the central nervous system host response to systemic lipopolysaccharide administration. Journal of Neuroinflammation 8, 124.

47. Ito, H.T., Smith, S.E.P., Hsiao, E., and Patterson, P.H. (2010). Maternal Immune Activation Alters Nonspatial Information Processing in the Hippocampus of the Adult Offspring. Brain Behav. Immun. 24, 930–941.

48. Jiang, H.-Y., Xu, L.-l., Shao, L., Xia, R.-M., Yu, Z.-H., Ling, Z.-X., Yang, F., Deng, M., and Ruan, B. (2016). Maternal infection during pregnancy and risk of autism spectrum disorders: A systematic review and meta-analysis. Brain Behav. Immun. 58, 165–172.

49. Jin, M., Jang, E., and Suk, K. (2014). Lipocalin-2 Acts as a Neuroinflammatogen in Lipopolysaccharide-injected Mice. Exp Neurobiol 23, 155–162.

50. Jongbloets, B.C., Lemstra, S., Schellino, R., Broekhoven, M.H., Parkash, J., Hellemons, A.J., Mao, T., Giacobini, P., van Praag, H., De Marchis, S., et al. (2017). Stage-specific functions of Semaphorin7A during adult hippocampal neurogenesis rely on distinct receptors. Nat Commun 8, 14666.

51. Kang, S.S., Ren, Y., Liu, C.C., Kurti, A., Baker, K.E., Bu, G., Asmann, Y., and Fryer, J.D. (2018). Lipocalin-2 protects the brain during inflammatory conditions. Molecular Psychiatry 23, 344–350.

52. Kentner, A.C., Bilbo, S.D., Brown, A.S., Hsiao, E.Y., McAllister, A.K., Meyer, U., Pearce, B.D., Pletnikov, M.V., Yolken, R.H., and Bauman, M.D. (2019). Maternal immune activation: reporting guidelines to improve the rigor, reproducibility, and transparency of the model. Neuropsychopharmacology 44, 245–258.

53. Khalaf-Nazzal, R., and Francis, F. (2013). Hippocampal development – Old and new findings. Neuroscience 248, 225–242.

54. Kim, J.-H., Kang, R.J., Hyeon, S.J., Ryu, H., Joo, H., Bu, Y., Kim, J.-H., and Suk, K. (2023). Lipocalin-2 Is a Key Regulator of Neuroinflammation in Secondary Traumatic and Ischemic Brain Injury. Neurotherapeutics 20, 803–821.

55. Kim, J.-H., Michiko, N., Choi, I.-S., Kim, Y., Jeong, J.-Y., Lee, M.-G., Jang, I.-S., and Suk, K. (2024). Aberrant activation of hippocampal astrocytes causes neuroinflammation and cognitive decline in mice. PLOS Biology 22, e3002687.

56. Knapska, E., Lioudyno, V., Kiryk, A., Mikosz, M., Górkiewicz, T., Michaluk, P., Gawlak, M., Chaturvedi, M., Mochol, G., Balcerzyk, M., et al. (2013). Reward learning requires activity of matrix metalloproteinase-9 in the central amygdala. J. Neurosci. 33, 14591–14600.

57. Knapska, E., Walasek, G., Nikolaev, E., Neuhäusser-Wespy, F., Lipp, H.-P., Kaczmarek, L., and Werka, T. (2006). Differential involvement of the central amygdala in appetitive versus aversive learning. Learning & Memory (Cold Spring Harbor, N.Y.) 13, 192–200.

58. Kuzniewska, B., Nader, K., Dabrowski, M., Kaczmarek, L., and Kalita, K. (2016). Adult Deletion of SRF Increases Epileptogenesis and Decreases Activity-Induced Gene Expression. Molecular neurobiology 53, 1478–1493.

59. Lee, S., Kim, J.-H., Kim, J.-H., Seo, J.-W., Han, H.-S., Lee, W.-H., Mori, K., Nakao, K., Barasch, J., and Suk, K. (2011). Lipocalin-2 Is a Chemokine Inducer in the Central Nervous System. The Journal of Biological Chemistry 286, 43855–43870.

60. Lee, S., Park, J.-Y., Lee, W.-H., Kim, H., Park, H.-C., Mori, K., and Suk, K. (2009). Lipocalin-2 Is an Autocrine Mediator of Reactive Astrocytosis. J. Neurosci. 29, 234–249.

61. Leekam, S.R., Prior, M.R., and Uljarevic, M. (2011). Restricted and repetitive behaviors in autism spectrum disorders: A review of research in the last decade. Psychological Bulletin 137, 562–593.

62. Legutko, D., Kuzniewska, B., Kalita, K., Yasuda, R., Kaczmarek, L., and Michaluk, P. (2024). BDNF signaling requires Matrix Metalloproteinase-9 during structural synaptic plasticity. bioRxiv.

63. Lepeta, K., Purzycka, K.J., Pachulska-Wieczorek, K., Mitjans, M., Begemann, M., Vafadari, B., Bijata, K., Adamiak, R.W., Ehrenreich, H., Dziembowska, M., et al. (2017). A normal genetic variation modulates synaptic MMP-9 protein levels and the severity of schizophrenia symptoms. EMBO Mol Med 9, 1100–1116.

64. Li, J., Xu, P., Hong, Y., Xie, Y., Peng, M., Sun, R., Guo, H., Zhang, X., Zhu, W., Wang, J., et al. (2023). Lipocalin-2-mediated astrocyte pyroptosis promotes neuroinflammatory injury via NLRP3 inflammasome activation in cerebral ischemia/reperfusion injury. Journal of Neuroinflammation 20, 148.

65. Li, M., Cui, Z., Niu, Y., Liu, B., Fan, W., Yu, D., and Deng, J. (2010). Synaptogenesis in the developing mouse visual cortex. Brain Research Bulletin 81, 107–113.

66. Liu, R., Wang, J., Chen, Y., Collier, J.M., Capuk, O., Jin, S., Sun, M., Mondal, S.K., Whiteside, T.L., Stolz, D.B., et al. (2022). NOX activation in reactive astrocytes regulates astrocytic LCN2 expression and neurodegeneration. Cell Death Dis 13, 1–15.

67. Lombardo, M.V., Moon, H.M., Su, J., Palmer, T.D., Courchesne, E., and Pramparo, T. (2018). Maternal immune activation dysregulation of the fetal brain transcriptome and relevance to the pathophysiology of autism spectrum disorder. Molecular Psychiatry 23, 1001–1013.

68. Lv, N., Wang, Y., Zhao, M., Dong, L., and Wei, H. (2021). The Role of PAX2 in Neurodevelopment and Disease. Neuropsychiatr Dis Treat 17, 3559–3567.

69. Malkova, N.V., Yu, C.Z., Hsiao, E.Y., Moore, M.J., and Patterson, P.H. (2012). Maternal immune activation yields offspring displaying mouse versions of the three core symptoms of autism. Brain Behav. Immun. 26, 607–616.

70. Marques, F., Rodrigues, A.-J., Sousa, J.C., Coppola, G., Geschwind, D.H., Sousa, N., Correia-Neves, M., and Palha, J.A. (2008). Lipocalin 2 is a Choroid Plexus Acute-Phase Protein. J Cereb Blood Flow Metab 28, 450–455.

71. Mattei, D., Djodari-Irani, A., Hadar, R., Pelz, A., de Cossío, L.F., Goetz, T., Matyash, M., Kettenmann, H., Winter, C., and Wolf, S.A. (2014). Minocycline rescues decrease in neurogenesis, increase in microglia cytokines and deficits in sensorimotor gating in an animal model of schizophrenia. Brain Behav. Immun. 38, 175–184.

72. Meyer, U., Feldon, J., and Fatemi, S.H. (2009). In-vivo rodent models for the experimental investigation of prenatal immune activation effects in neurodevelopmental brain disorders. Neuroscience and Biobehavioral Reviews 33, 1061–1079.

73. Michaluk, P., Wawrzyniak, M., Alot, P., Szczot, M., Wyrembek, P., Mercik, K., Medvedev, N., Wilczek, E., De Roo, M., Zuschratter, W., et al. (2011). Influence of matrix metalloproteinase MMP-9 on dendritic spine morphology. J Cell Sci 124, 3369–3380.

74. Mucha, M., Skrzypiec, A.E., Schiavon, E., Attwood, B.K., Kucerova, E., and Pawlak, R. (2011). Lipocalin-2 controls neuronal excitability and anxiety by regulating dendritic spine formation and maturation. Proceedings of the National Academy of Sciences of the United States of America 108, 18436–18441.

75. Nabeel Mustafa, A., Salih Mahdi, M., Ballal, S., Chahar, M., Verma, R., Ali Al-Nuaimi, A.M., Kumar, M.R., Kadhim, A.A.-H.R., Adil, M., and Jasem Jawad, M. (2025). Netrin-1: Key insights in neural development and disorders. Tissue Cell 93, 102678.

76. Nouel, D., Burt, M., Zhang, Y., Harvey, L., and Boksa, P. (2012). Prenatal exposure to bacterial endotoxin reduces the number of GAD67-and reelin-immunoreactive neurons in the hippocampus of rat offspring. Eur Neuropsychopharmacol 22, 300–307.

77. Oskvig, D.B., Elkahloun, A.G., Johnson, K.R., Phillips, T.M., and Herkenham, M. (2012). Maternal immune activation by LPS selectively alters specific gene expression profiles of interneuron migration and oxidative stress in the fetus without triggering a fetal immune response. Brain Behav Immun 26, 623–634.

78. Pasciuto, E., Borrie, S.C., Kanellopoulos, A.K., Santos, A.R., Cappuyns, E., D’Andrea, L., Pacini, L., and Bagni, C. (2015). Autism Spectrum Disorders: Translating human deficits into mouse behavior. Neurobiol Learn Mem 124, 71–87.

79. Pendyala, G., Chou, S., Jung, Y., Coiro, P., Spartz, E., Padmashri, R., Li, M., and Dunaevsky, A. (2017). Maternal Immune Activation Causes Behavioral Impairments and Altered Cerebellar Cytokine and Synaptic Protein Expression. Neuropsychopharmacology 42, 1435–1446.

80. Provenzano, G., Gilardoni, A., Maggia, M., Pernigo, M., Sgado, P., Casarosa, S., and Bozzi, Y. (2020). Altered Expression of GABAergic Markers in the Forebrain of Young and Adult Engrailed-2 Knockout Mice. Genes (Basel) 11.

81. Punetha, J., Karaca, E., Gezdirici, A., Lamont, R.E., Pehlivan, D., Marafi, D., Appendino, J.P., Hunter, J.V., Akdemir, Z.C., Fatih, J.M., et al. (2019). Biallelic CACNA2D2 variants in epileptic encephalopathy and cerebellar atrophy. Ann Clin Transl Neurol 6, 1395–1406.

82. Puścian, A., Łęski, S., Kasprowicz, G., Winiarski, M., Borowska, J., Nikolaev, T., Boguszewski, P.M., Lipp, H.-P., and Knapska, E. (2016). Eco-HAB as a fully automated and ecologically relevant assessment of social impairments in mouse models of autism. eLife 5, e19532.

83. Puścian, A., Winiarski, M., Borowska, J., Łęski, S., Górkiewicz, T., Chaturvedi, M., Nowicka, K., Wołyniak, M., Chmielewska, J.J., Nikolaev, T., et al. (2022). Targeted therapy of cognitive deficits in fragile X syndrome. Molecular Psychiatry 27, 2766–2776.

84. Rossanti, R., Morisada, N., Nozu, K., Kamei, K., Horinouchi, T., Yamamura, T., Minamikawa, S., Fujimura, J., Nagano, C., Sakakibara, N., et al. (2020). Clinical and genetic variability of PAX2-related disorder in the Japanese population. J Hum Genet 65, 541–549.

85. Roszkowska, M., Krysiak, A., Majchrowicz, L., Nader, K., Beroun, A., Michaluk, P., Pekala, M., Jaworski, J., Kondrakiewicz, L., Puścian, A., et al. (2022). SRF depletion in early life contributes to social interaction deficits in the adulthood. Cellular and molecular life sciences: CMLS 79, 278.

86. Rydzanicz, M., Kuzniewska, B., Magnowska, M., Wojtowicz, T., Stawikowska, A., Hojka, A., Borsuk, E., Meyza, K., Gewartowska, O., Gruchota, J., et al. (2024). Mutation in the mitochondrial chaperone TRAP1 leads to autism with more severe symptoms in males. EMBO Mol Med 16, 2976–3004.

87. Saatci, D., van Nieuwenhuizen, A., and Handunnetthi, L. (2021). Maternal infection in gestation increases the risk of non-affective psychosis in offspring: a meta-analysis. J Psychiatr Res 139, 125–131.

88. Sen, B., Singh, A.S., Sinha, S., Chatterjee, A., Ahmed, S., Ghosh, S., and Usha, R. (2010). Family-based studies indicate association of Engrailed 2 gene with autism in an Indian population. Genes Brain Behav 9, 248–255.

89. Serafini, T., Colamarino, S.A., Leonardo, E.D., Wang, H., Beddington, R., Skarnes, W.C., and Tessier-Lavigne, M. (1996). Netrin-1 is required for commissural axon guidance in the developing vertebrate nervous system. Cell 87, 1001–1014.

90. Sgado, P., Genovesi, S., Kalinovsky, A., Zunino, G., Macchi, F., Allegra, M., Murenu, E., Provenzano, G., Tripathi, P.P., Casarosa, S., et al. (2013). Loss of GABAergic neurons in the hippocampus and cerebral cortex of Engrailed-2 null mutant mice: implications for autism spectrum disorders. Exp Neurol 247, 496–505.

91. Shi, L., Smith, S.E.P., Malkova, N., Tse, D., Su, Y., and Patterson, P.H. (2009). Activation of the maternal immune system alters cerebellar development in the offspring. Brain Behav. Immun. 23, 116–123.

92. Shin, H.J., Jeong, E.A., Lee, J.Y., An, H.S., Jang, H.M., Ahn, Y.J., Lee, J., Kim, K.E., and Roh, G.S. (2021). Lipocalin-2 Deficiency Reduces Oxidative Stress and Neuroinflammation and Results in Attenuation of Kainic Acid-Induced Hippocampal Cell Death. Antioxidants 10, 100.

93. Skrzypiec, A.E., Shah, R.S., Schiavon, E., Baker, E., Skene, N., Pawlak, R., and Mucha, M. (2013). Stress-induced lipocalin-2 controls dendritic spine formation and neuronal activity in the amygdala. PLoS One 8, e61046.

94. Solek, C.M., Farooqi, N., Verly, M., Lim, T.K., and Ruthazer, E.S. (2018). Maternal immune activation in neurodevelopmental disorders. Developmental Dynamics 247, 588–619.

95. Souza, B.R., Codo, B.C., Romano-Silva, M.A., and Tropepe, V. (2024). Darpp-32 is regulated by dopamine and is required for the formation of GABAergic neurons in the developing telencephalon. Prog Neuropsychopharmacol Biol Psychiatry 134, 111060.

96. Sugihara, G., Hashimoto, K., Iwata, Y., Nakamura, K., Tsujii, M., Tsuchiya, K.J., Sekine, Y., Suzuki, K., Suda, S., Matsuzaki, H., et al. (2007). Decreased serum levels of hepatocyte growth factor in male adults with high-functioning autism. Prog Neuropsychopharmacol Biol Psychiatry 31, 412–415.

97. Szepesi, Z., Bijata, M., Ruszczycki, B., Kaczmarek, L., and Wlodarczyk, J. (2013). Matrix metalloproteinases regulate the formation of dendritic spine head protrusions during chemically induced long-term potentiation. PLoS One 8, e63314.

98. Szewczyk, L.M., Lipiec, M.A., Liszewska, E., Meyza, K., Urban-Ciecko, J., Kondrakiewicz, L., Goncerzewicz, A., Rafalko, K., Krawczyk, T.G., Bogaj, K., et al. (2024). Astrocytic β-catenin signaling via TCF7L2 regulates synapse development and social behavior. Molecular Psychiatry 29, 57–73.

99. Tajima, Y., Ito, K., Yuan, Y., Frank, M.O., Saito, Y., and Darnell, R.B. (2023). NOVA1 acts on Impact to regulate hypothalamic function and translation in inhibitory neurons. Cell Rep 42, 112050.

100. Thomas, A., Burant, A., Bui, N., Graham, D., Yuva-Paylor, L.A., and Paylor, R. (2009). Marble burying reflects a repetitive and perseverative behavior more than novelty-induced anxiety. Psychopharmacology 204, 361–373.

101. Tioleco, N., Silberman, A.E., Stratigos, K., Banerjee-Basu, S., Spann, M.N., Whitaker, A.H., and Turner, J.B. (2021). Prenatal maternal infection and risk for autism in offspring: A meta-analysis. Autism Research 14, 1296–1316.

102. Tunster, S.J. (2017). Genetic sex determination of mice by simplex PCR. Biol Sex Differ 8, 31.

103. Vichaya, E.G., Gross, P.S., Estrada, D.J., Cole, S.W., Grossberg, A.J., Evans, S.E., Tuvim, M.J., Dickey, B.F., and Dantzer, R. (2019). Lipocalin-2 is dispensable in inflammation-induced sickness and depression-like behavior. Psychopharmacology (Berl) 236, 2975–2982.

104. Wischhof, L., Irrsack, E., Osorio, C., and Koch, M. (2015). Prenatal LPS-exposure--a neurodevelopmental rat model of schizophrenia--differentially affects cognitive functions, myelination and parvalbumin expression in male and female offspring. Prog Neuropsychopharmacol Biol Psychiatry 57, 17–30.

105. Wu, Y., Qi, F., Song, D., He, Z., Zuo, Z., Yang, Y., Liu, Q., Hu, S., Wang, X., Zheng, X., et al. (2018). Prenatal influenza vaccination rescues impairments of social behavior and lamination in a mouse model of autism. Journal of Neuroinflammation 15, 228.

106. Xing, C., Wang, X., Cheng, C., Montaner, J., Mandeville, E., Leung, W., van Leyen, K., Lok, J., Wang, X., and Lo, E.H. (2014). Neuronal Production of Lipocalin-2 as a Help-Me Signal for Glial Activation. Stroke 45, 2085–2092.

107. Yan, L., Yang, F., Wang, Y., Shi, L., Wang, M., Yang, D., Wang, W., Jia, Y., So, K.-F., and Zhang, L. (2024). Stress increases hepatic release of lipocalin 2 which contributes to anxiety-like behavior in mice. Nature Communications 15, 3034.

108. Zhang, C., Frias, M.A., Mele, A., Ruggiu, M., Eom, T., Marney, C.B., Wang, H., Licatalosi, D.D., Fak, J.J., and Darnell, R.B. (2010). Integrative modeling defines the Nova splicing-regulatory network and its combinatorial controls. Science 329, 439–443.

109. Zhang, P.-X., Zhang, F.-R., Xie, J.-J., Tao, L.-H., Lü, Z., Xu, X.-E., Shen, J., Xu, L.-Y., and Li, E.-M. (2012). Expression of NGAL and NGALR in human embryonic, fetal and normal adult tissues. Molecular Medicine Reports 6, 716–722.

110. Zhang, Z., and van Praag, H. (2015). Maternal immune activation differentially impacts mature and adult-born hippocampal neurons in male mice. Brain Behav. Immun. 45, 60–70.

111. Zhao, J., Chen, H., Zhang, M., Zhang, Y., Qian, C., Liu, Y., He, S., Zou, Y., and Liu, H. (2016). Early expression of serum neutrophil gelatinase-associated lipocalin (NGAL) is associated with neurological severity immediately after traumatic brain injury. Journal of the Neurological Sciences 368, 392–398.

112. Zhou, Y.-y., Zhang, W.-w., Chen, F., Hu, S.-s., and Jiang, H.-y. (2021). Maternal infection exposure and the risk of psychosis in the offspring: A systematic review and meta-analysis. J Psychiatr Res 135, 28–36.

113. Zhou, Y., Zhou, B., Pache, L., Chang, M., Khodabakhshi, A.H., Tanaseichuk, O., Benner, C., and Chanda, S.K. (2019). Metascape provides a biologist-oriented resource for the analysis of systems-level datasets. Nat Commun 10, 1523.

114. Zhu, C.-Y., Jiang, H.-Y., and Sun, J.-J. (2022). Maternal infection during pregnancy and the risk of attention-deficit/hyperactivity disorder in the offspring: A systematic review and meta-analysis. Asian J Psychiatr 68, 102972.

